# Processing of Pain by the Developing Brain: Evidence of Differences Between Adolescent and Adult Females

**DOI:** 10.1101/2021.05.14.444209

**Authors:** Han Tong, Thomas C. Maloney, Michael F. Payne, Christopher D. King, Tracy V. Ting, Susmita Kashikar-Zuck, Robert C. Coghill, Marina López-Solà

## Abstract

Adolescence is a sensitive period for both brain development and the emergence of chronic pain particularly in females. However, the brain mechanisms supporting pain perception during adolescence remain unclear. This study compares perceptual and brain responses to pain in female adolescents and adults to characterize pain processing in the developing brain. Thirty adolescent (ages 13-17) and thirty adult (ages 35-55) females underwent a functional MRI scan involving acute experimental pain. Participants received 12 ten-second noxious pressure stimuli which were applied to the left thumbnail at 2.5 and 4 kg/cm^2^, and rated pain intensity and unpleasantness on a visual analogue scale. We found a significant group-by-stimulus intensity interaction on pain ratings. Compared to adults, adolescents reported greater pain intensity and unpleasantness in response to 2.5 kg/cm^2^, but not 4 kg/cm^2^. Adolescents showed greater medial-lateral prefrontal cortex (PFC) and supramarginal gyrus activation in response to 2.5 kg/cm^2^, and greater medial PFC and rostral anterior cingulate responses to 4 kg/cm^2^. Adolescents showed augmented pain-evoked responses in the Neurologic Pain Signature and greater activation in the default mode (DMN) and ventral attention (VAN) networks. Also, the amygdala and associated regions played a stronger role in predicting pain intensity in adolescents, and activity in DMN and VAN regions more strongly mediated the relationship between stimulus intensity and pain ratings. This study provides the first evidence of augmented pain-evoked brain responses in healthy female adolescents involving regions important for nociceptive, affective and cognitive processing, in line with their augmented sensitivity to low-intensity noxious stimuli.

## 1. Introduction

Pain is a major health issue that plagues adolescence. Studies have found that 20-46% of adolescents worldwide suffer from chronic weekly pain[30,44,71]. Indeed, adolescence marks a time when gender differences emerge and significant increases in the prevalence of chronic pain conditions are seen in adolescent females[44,52,71], many of which persist into adulthood, such as fibromyalgia[43, 100], complex regional pain syndrome[1] and irritable bowel syndrome[38]. Their emergence at this stage of development raises interesting questions about what specific changes related to pain processing occur during puberty that make adolescent females more vulnerable. Although the past two decades have seen a great advancement in our understanding of pain in adults[2,14,17,94], little is known about characteristics of pain processing in adolescents. To our knowledge, no study has directly compared pain sensitivity and brain responses to pain between adolescents and adults. Previous studies have shown that pain sensitivity generally decreases with age[23]. One study found a rapid rise in cutaneous pain threshold to the age of 25[92]. This observed greater pain sensitivity during development may involve peripheral and central nervous system mechanisms. On the one hand, adolescents have a higher density of intra-epidermal nerve fibers (i.e., unmyelinated nociceptors), suggesting potentially increased nociceptive input to the central nervous system[53, 70]. On the other, adolescence is a critical period for brain development when the brain undergoes a fundamental reorganization[74], permitting various environmental influences to exert powerful effects that could determine health and social outcomes in adulthood[22, 47]. In particular, significant morphological and functional changes occur in amygdala and associated regions during adolescence[34,35,37,66], which may account for heightened emotional reactivity to aversive stimuli[12, 88]. Furthermore, association cortices such as the prefrontal cortex (PFC) and the posterior parietal cortex (PPC), which contribute greatly to forming and regulating pain experience[2,10,59], undergo continued structural and functional maturation during adolescence[12, 31]. Moreover, the default mode network (DMN), another key player in pain perception and regulation in both health and disease[6,7,50,55,63,90], undergoes maturation during adolescence by increasing intra-network integration and inter-network segregation[24, 83]. In this study, we compared psychophysical and brain responses to controlled noxious pressure stimulations between adolescents and middle-aged adults. We only enrolled female participants because most primary chronic pain conditions of adolescence predominantly affect females[26, 93], and there could be qualitative sex differences in pain processing which would need to be examined separately[64, 65]. We sought to identify the neural processes in the brain that characterize adolescents’ pain experience. To this end, besides standard univariate analyses, we conducted whole-brain multilevel mediation analyses and computed pain-evoked responses in large-scale cortical networks[99] and the Neurologic Pain Signature (NPS)[94]. The NPS was used as a summary measure of nociceptive-specific signal processing at the brain level since it is particularly sensitive to nociception-dependent physical pain, but not other aversive experiences[48,56,57,58,60,94,95]. We expected that, compared to adults, adolescents would show: (1) greater pain sensitivity accompanied by augmented pain-evoked nociceptive-specific NPS responses (in agreement with greater nociceptive fiber afference), and (2) augmented responses in brain regions involved in regulating emotional responses and cognitive appraisal of painful aversive stimuli, such as the amygdala and related regions, the medial and lateral aspects of the prefrontal cortex and the DMN, all also undergoing maturation during adolescence.

## 2. Materials and methods

### 2.1. Participants

This study included 30 healthy adolescent girls (13-17 years old, mean age of 16.00 ± 1.25 years) and 30 healthy women (35-55 years old, mean age of 44.67 ± 6.29 years) without acute pain (assessed by the 0-10 numeric pain rating scale) and any history of psychiatric, neurological, or chronic pain disorders. Before being enrolled in the study, all adult participants and the parents of the adolescent participants provided written informed consent. In addition, all adolescent participants provided informed assent. The study protocol and consent forms were approved by Cincinnati Children’s Hospital Medical Center Institutional Review Board (Study ID: 2017-7771). All participants completed the functional magnetic resonance imaging (fMRI) task and received compensation for their participation. All of the data needed for this study were collected between February 2018 and December 2019 and used for subsequent analyses.

### 2.2 Study procedures

This study consisted of two sessions. Session 1 involved collecting demographic and biometric information and familiarizing the participants with the pressure stimulation device and the pressure pain task. Session 2 immediately followed session 1 and involved functional and anatomical brain MRI scans.

#### 2.2.1 Pressure pain device

As in previous studies[32,59,60,78], a calibrated computer-controlled pneumatic device, which can reliably transmit preset pressure to 1-cm^2^ surface, was used to deliver noxious pressure stimuli to the base of the participants’ left thumbnail. Two pressure levels were applied in this experiment, a low stimulus intensity of 2.5 kg/cm^2^ and a moderate stimulus intensity of 4 kg/cm^2^.

#### 2.2.2 Noxious pressure stimulation fMRI task

We adopted a block-design for our noxious pressure stimulation fMRI task, programmed and presented to the participants using the E-Prime 3.0 software (Psychology Software Tools, Pittsburgh, PA). As shown in **Figure 1**, this task was composed of two consecutive fMRI runs (i.e., scanning sequences), each containing 6 trials (three at each pressure level, in a mixed pseudorandom order). Each trial began with a rest period with pseudorandom duration (range: 11-20 seconds), followed by a brief auditory stimulus (200-millisecond tone), a 3-to-6-second anticipatory period, and then a fixed 10-second pressure stimulation period. After an 8-10 second post-stimulation rest period, the participants were asked to rate pain intensity (“How intense was the pain you just experienced?”) and pain unpleasantness (“How unpleasant was the pain you just experienced?”) on computerized visual analogue scales from 0 (not painful/unpleasant at all) to 100 (most painful/unpleasant imaginable)[75, 76]. The participants were instructed to move the cursor on the scales using an MRI-compatible trackball until the position that best describes their pain experience and click the button to submit their ratings. The numbers between 0 and 100 on the scales were not visible to the participants.

**Figure 1.**
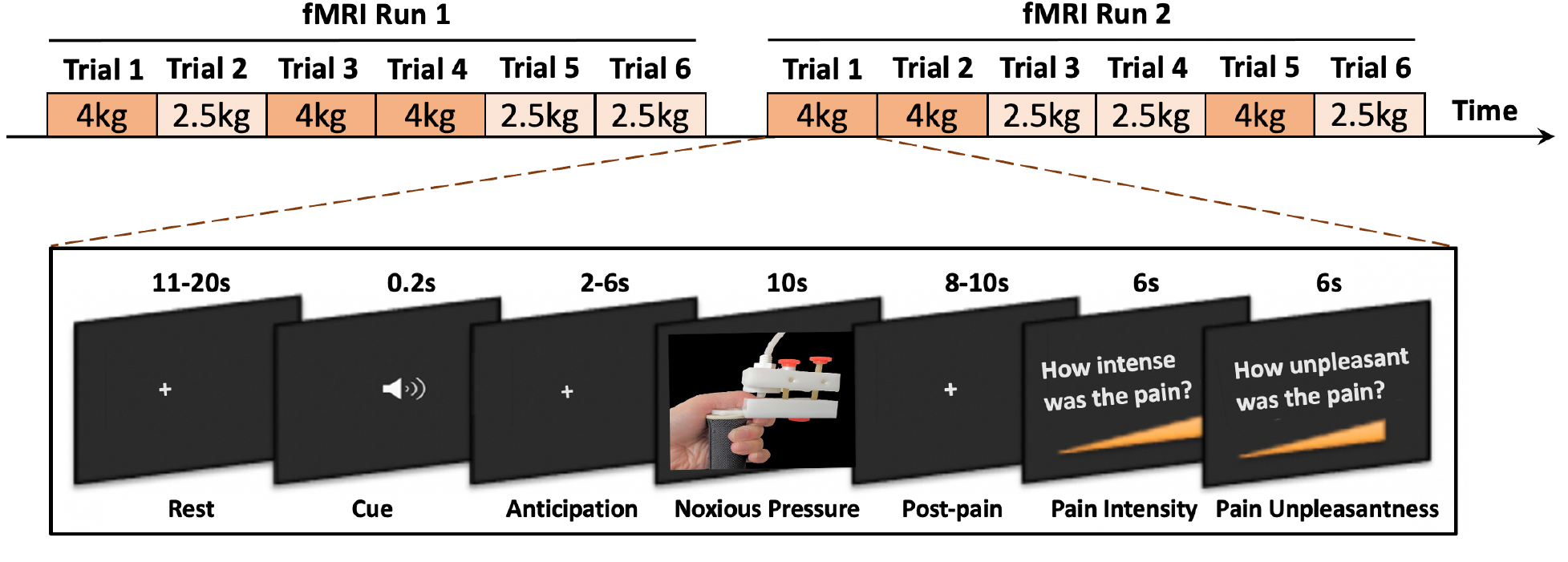
Graphic representation of the noxious pressure stimulation fMRI task.

### 2.3 Magnetic resonance imaging data acquisition

All MRI data for this study was acquired using a Philips Ingenia 3.0T MR System (Philips Healthcare, Best, The Netherlands) with a 32-channel head coil at Cincinnati Children’s Hospital Medical Center. Structural images of the brain were acquired using the standard T1 weighted gradient echo sequence with the following scan parameters: TR = 10 ms, TE = 1.8, 3.8, 5.8, 7.8 ms, field of view = 256 x 224 x 200 mm, voxel size = 1 x 1 x 1 mm, number of slices = 200, flip angle = 8°, slice orientation = sagittal, and total scan duration = 4:42 minutes. Blood oxygen level-dependent (BOLD) fMRI data were collected using T2* weighted echo planar imaging sequence with multiband sensitivity encoding (SENSE) technique[25,51,77]. Scan parameters for the BOLD fMRI acquisition were as follows: multiband acceleration factor = 4, TR = 650 ms, TE =30 ms, field of view = 200 mm, flip angle = 53°, voxel size =2.5 x 2.5 x 3.5 mm, slice orientation = transverse (parallel to the orbitofrontal cortex line), slice thickness = 3.5 mm, number of slice = 40 (provided whole-brain coverage), number of volumes = 522, dummy scans = 12, and total scan duration = 5:42 minutes.

### 2.4 Data analyses

#### 2.4.1 Statistical analyses of behavioral data

Mixed-design ANOVA with “group” as a between-subject variable and “pressure” as a within-subject variable was performed using R software (version 3.6.2, R Foundation for Statistical Computing, Vienna, Austria) to assess differences in pain intensity and unpleasantness under two experimental conditions (i.e., noxious pressure stimuli at 2.5kg/cm^2^ and 4kg/cm^2^) between the adolescent group and the adult group. Post-hoc between-group comparisons for each experimental condition were made using Fisher’s least significant difference (LSD) method.

#### 2.4.2 Preprocessing of neuroimaging data

The neuroimaging data was preprocessed using FSL (FMRIB Software Library version 6.0.3, the Analysis Group, FMRIB, Oxford, UK)[40, 87] and AFNI (Analysis of Functional Neuroimages version 20.3.02, Medical College of Wisconsin, WI, USA)[20]. For the T1-weighted structural image of each participant, brain extraction was performed using FSL’s BET (Brain Extraction Tool)[86], then bias correction and segmentation were carried out using FSL’s FAST (FMRIB’s Automated Segmentation Tool)[103]. The brain extracted image was then normalized and resampled to the 2-mm isotropic MNI ICBM 152 non-linear 6th generation template[27] using FSL’s FLIRT (FMRIB’s Linear Image Registration Tool)[39, 41]. Each participant’s functional (BOLD) scans were preprocessed in the following steps: First, brain extraction was carried out using FSL’s BET[86]. Next, outlying functional volumes (i.e. spikes) were detected using the DVARS metric within FSL’s “fsl_motion_outliers”[73]. Motion correction of the BOLD time-series was carried out using MCFLIRT[39]. The motion corrected data was high-pass filtered at 0.01 Hz (100 seconds) and smoothed with a 6 mm FWHM filter using AFNI’s 3dBandpass. Intensity normalization (i.e., scaling each functional volume by its mean global intensity) was applied to minimize confounds arising from pain-induced global CBF fluctuations[15,16,101,102]. The intensity-normalized data were then aligned to the MNI template[27] by first co-registering it with the participant’s T1 structural MPRAGE image using FSL’s FLIRT (6-parameter rigid body model)[39, 41].

#### 2.4.3 First-level general linear model analyses

We modeled each run of the preprocessed functional MRI data for each participant using the general linear model (GLM) approach as implemented in FSL’s “fsl_glm”[97] to estimate each participant’s brain responses to pain in the following two ways: (1) modeling the three pain periods associated with 2.5 kg/cm^2^ stimuli as one regressor and the other three pain periods associated with 4 kg/cm^2^ stimuli as another regressor to prepare the data for higher-level GLM analyses and Neurologic Pain Signature (NPS) analyses; (2) modeling each of the six pain periods as a separate regressor to be used in the whole-brain multilevel mediation analyses. In addition to the pain period regressors, our GLM model included regressors for the anticipatory periods, post-pain periods, and pain rating periods. The remaining “rest” period was used as the implicit baseline. Finally, six motion parameters (three for translational motion and three for rotational motion) and outlying volumes (spikes) were included as nuisance regressors (**Figure S1**).

#### 2.4.4 Higher-level general linear model analyses

The two runs of each participant’s first-level GLM results, which included estimated contrasts of parameter estimates (COPEs) and their variances (VARCOPEs), were combined at the second level (single-subject level) using the fixed effects modeling in FSL with “flameo”[96]. Then at the third level (group-level), mixed effects modeling (FLAME 1+2)[96] was used to compute each group’s mean brain responses to pressure pain (one-sample t-test) and between-group differences (two-sample t-test) for each condition (2.5 kg/cm^2^ and 4 kg/cm^2^). The results of third-level analyses were corrected for multiple comparisons across the whole brain using FSL’s

“cluster” tool. Clusters of voxels were identified using a threshold of Z>3.1 and their statistical significance (p<0.05) was estimated by cluster-based inference according to Gaussian random field theory[98].

#### 2.4.5 Pain-evoked Neurologic Pain Signature responses

As a multivariate brain pattern that specifically responds to somatic pain rather than to other aversive experiences, the NPS was used to further investigate nociceptive-specific neural responses in adolescents and adults. A single scalar value summarizing each participant’s NPS signature response was computed for the two pressure pain conditions (i.e., 2.5 kg/cm^2^ and 4 kg/cm^2^) in each run respectively. Specifically, we computed the dot product of the voxel weights within the pre-defined NPS mask and the contrast image of parameter estimates from first-level GLM analyses for each subject and run using custom code developed in Python (version 3.7.4, Python Software Foundation, OR, USA) that utilizes the Nibabel[9] and Numpy[36] packages. Next, the NPS signature responses for the two runs were averaged for each participant. Last, a group by pressure mixed ANOVA was performed using R software (version 3.6.2, R Foundation for Statistical Computing, Vienna, Austria) to compare the mean NPS responses to noxious stimuli at 2.5kg/cm^2^ and 4kg/cm^2^ between the adolescent and adult groups. Post-hoc between-group comparisons for each stimulus intensity were made using Fisher’s least significant difference (LSD) method.

#### 2.4.6 Pain-evoked neural responses in large-scale brain networks

In order to assess how pain-evoked neural responses mapped onto large-scale functional brain networks, we computed the dot product, using our python code, of each participant’s contrast images of parameter estimates for each run (i.e., “pressure pain at 2.5 kg/cm^2^” and “pressure pain at 4 kg/cm^2^”) and pre-defined masks of the previously identified seven major cortical resting-state networks[99], including the somatomotor network, the default mode network, the fronto-parietal network, the limbic network, the ventral attentional network, the dorsal attentional network, the limbic network and the visual network. Then, the responses within each brain network for each run were combined by taking an arithmetic mean at the individual participant level, which resulted in a single-scalar value representing a summary metric of neural responses to pain across the entire functional brain network. Finally, between-group comparisons were carried out in R software for each network and each condition using two-sample t-tests.

#### 2.4.7 Whole-brain multilevel mediation analyses

First-level contrast images for the single-trial pain period regressors for each participant were carried forward to a multilevel mediation analysis model. To avoid that single-trial estimates could be driven by movement artifacts or other sources of noise, trial estimates with variance inflation factor of 5 or more were excluded from further analysis[45,56,58]. We then tested relationships between conditions (noxious stimulus intensity of 4 kg/cm^2^ vs 2.5 kg/cm^2^), single-trial pain-evoked brain responses (contrast images for each trial), and pain intensity ratings across individual trials using multilevel mediation analysis found in the Mediation Toolbox (canlab.github.io) and implemented in MATLAB (version R2019b, MathWorks, MA, USA)[4,5,45,56]. Multilevel mediation analysis identifies brain regions that show partially independent, but not orthogonal, effects: (1) brain regions that show activity increases or decreases during high vs. low painful stimulation (path a), (2) brain regions that predict changes in pain intensity (path b) even after controlling for path a, and (3) mediating regions (path a x b), i.e., regions most directly associated with both the experimental manipulation (high vs low painful stimulus) and variations in pain ratings. The idea underlying “mediation” is that painful stimulus intensity has an effect on pain perception that can be decomposed into 2 constituent pathways: painful stimulus intensity affects the brain response in some regions, which in turn leads to changes in pain perception. Some other regions that respond to stimulus intensity (path a) might not correlate with pain perception. In this case, they would not be mediators, because mediation requires both stimulus and pain effects (controlling for stimulus) to be present. Likewise, some areas that correlate with pain perception (path b) might not respond to stimulus intensity. These areas will also not appear as mediators. In this study, we were specifically interested in path b, showing activation increases that predict greater pain reports at the single trial level even after controlling for stimulus intensity, and path a x b of significant brain mediators of the effect of stimulus intensity on pain perception. The resulting activation maps were thresholded at q < 0.05 false discovery rate (FDR)-corrected within an extensive whole-brain gray-matter mask including 352,328 voxels, as previously done by our group and others[4,45,56,58]. To test the effect of group on the mediation paths of interest, we also added a second-level moderator (adolescents > adults) and the results of between-group comparisons were thresholded at p < 0.001[4, 45]. To facilitate interpretation of the functional maps, adjacent voxels to a corrected cluster were also displayed at lower thresholds of p<0.005 uncorrected.

## 3. Results

### 3.1 Adolescents have greater pain sensitivity than adults to low level of noxious pressure

**Figure 2** summarizes the pain ratings to noxious pressure stimuli by group and by pressure. The pain intensity and pain unpleasantness ratings to stimuli at 2.5 kg/cm^2^ were 22.71 ± 14.68 (mean ± std) and 20.92 ± 13.59 in adolescents, 13.75 ± 9.93 and 12.29 ± 11.45 in adults, respectively. The pain intensity and pain unpleasantness ratings to stimuli at 4 kg/cm^2^ were 31.64 ± 18.91 and 32.31 ± 19.39 in adolescents, 29.83 ± 17.32 and 27.79 ± 16.62, respectively. Mixed-design analysis of variance (ANOVA) with “group” as the between-subject factor and “pressure” as the within-subject factor was performed for pain intensity and pain unpleasantness ratings respectively. As expected, we found a significant main effect of pressure on pain intensity (F=92.09, p<0.0001) and pain unpleasantness ratings (F=95.42, p<0.0001), indicating that pain ratings increased with the rise of pressure level. We also observed a trend for a main effect of group on pain unpleasantness ratings (F=3.04, p=0.087) but not on pain intensity ratings (F=2.00, p=0.168). Moreover, we found a significant group × pressure interaction effect on pain intensity ratings (F=7.52, p=0.008), indicating that increases in pain ratings with rise in pressure level are different between adolescents and adults. The interaction effect was not significant for pain unpleasantness (F=2.23, p=0.141). Following ANOVA, we made post-hoc between-group comparisons for pain intensity and unpleasantness at each pressure level using Fisher’s least significant difference (LSD) method. Adolescent participants reported significantly greater pain intensity (t=2.23, p=0.030) and pain unpleasantness (t=2.15, p=0.036) than adult participants. Pain ratings in adolescents did not differ from adults in response to stimuli at 4 kg/cm^2^ (t=0.45, p=0.655 for pain intensity and t=1.12, p=0.265 for pain unpleasantness). These findings suggest that adolescents are more sensitive than adults to low level, peri-threshold noxious pressure stimuli.

**Figure 2.**
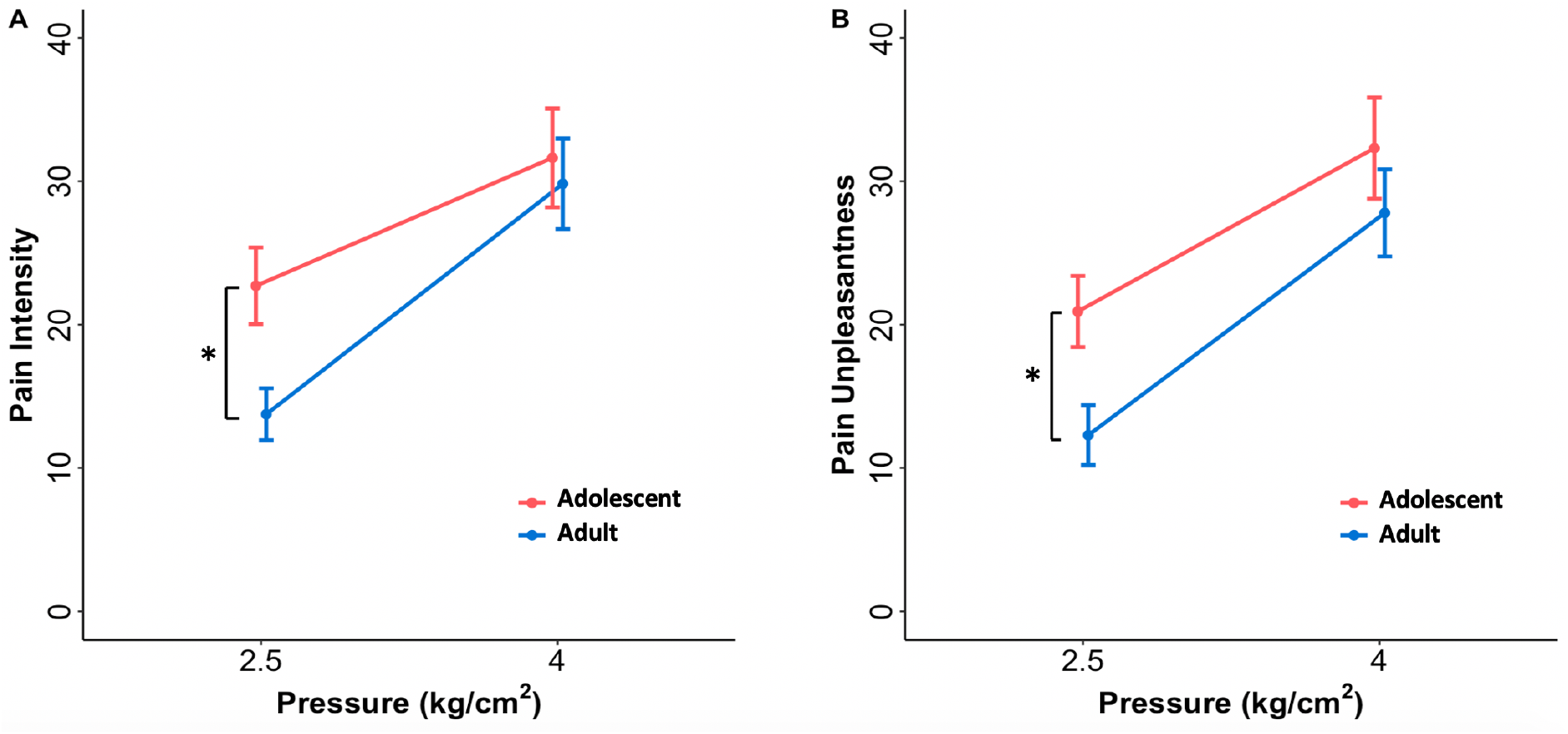
Pain intensity and pain unpleasantness ratings to noxious pressure stimuli by pressure and group. (A) Adolescents reported higher pain intensity than adults in response to noxious pressure stimuli at 2.5kg/cm2. However, this between-group difference disappeared at 4 kg/cm2. We found a significant main effect of pressure and an interaction effect of group by pressure on pain intensity ratings (see main text for statistics). (B) Adolescents reported higher pain unpleasantness than adults to noxious pressure stimuli at 2.5kg/cm2, but not to stimuli at 4kg/cm2. The group by pressure interaction effect on pain unpleasantness was not significant, but we found a significant main effect of pressure and a trend toward significant main effect of group (see main text for statistics). Error bars represent standard error of the mean. *p<0.05 in post-hoc t-test following mixed-design ANOVA.

### 3.2 Characterization of brain responses to pain in adolescents

#### 3.2.1 Adolescents exhibit greater pain-evoked neural responses than adults

Pain-evoked brain responses in adolescents involved brain regions similar to those found in adults (**Figure 3, Table S1-S4**), including bilateral insula/central operculum, anterior cingulate cortex, parietal operculum (S2), supramarginal gyrus, primary sensorimotor cortex (S1/M1), supplementary motor area, dorsolateral prefrontal cortex, superior temporal gyrus, basal ganglia, thalamus, periaqueductal gray matter and amygdala. Pain-evoked deactivations were found in the cerebellum, fusiform gyrus, precuneus/ posterior cingulate cortex, and occipital visual cortex. Additionally, adolescents showed significant pain-evoked activation in medial prefrontal cortex and deactivation in the medial orbitofrontal cortex, which were not found in adults. When statistically compared, adolescents exhibited significantly greater activation than adults in the dorsolateral prefrontal cortex, the dorsomedial prefrontal cortex, and supramarginal gyrus, along with greater deactivation in the medial orbitofrontal cortex, in response to noxious pressure stimuli at 2.5 kg/cm^2^ (**Figure 3, Table S5**). In response to noxious pressure stimuli at 4 kg/cm^2^, adolescents showed greater activations in rostral anterior cingulate and dorsomedial prefrontal cortex, along with greater deactivations in the cerebellum and fusiform gyrus (**Figure 3, Table S6**).

**Figure 3.**
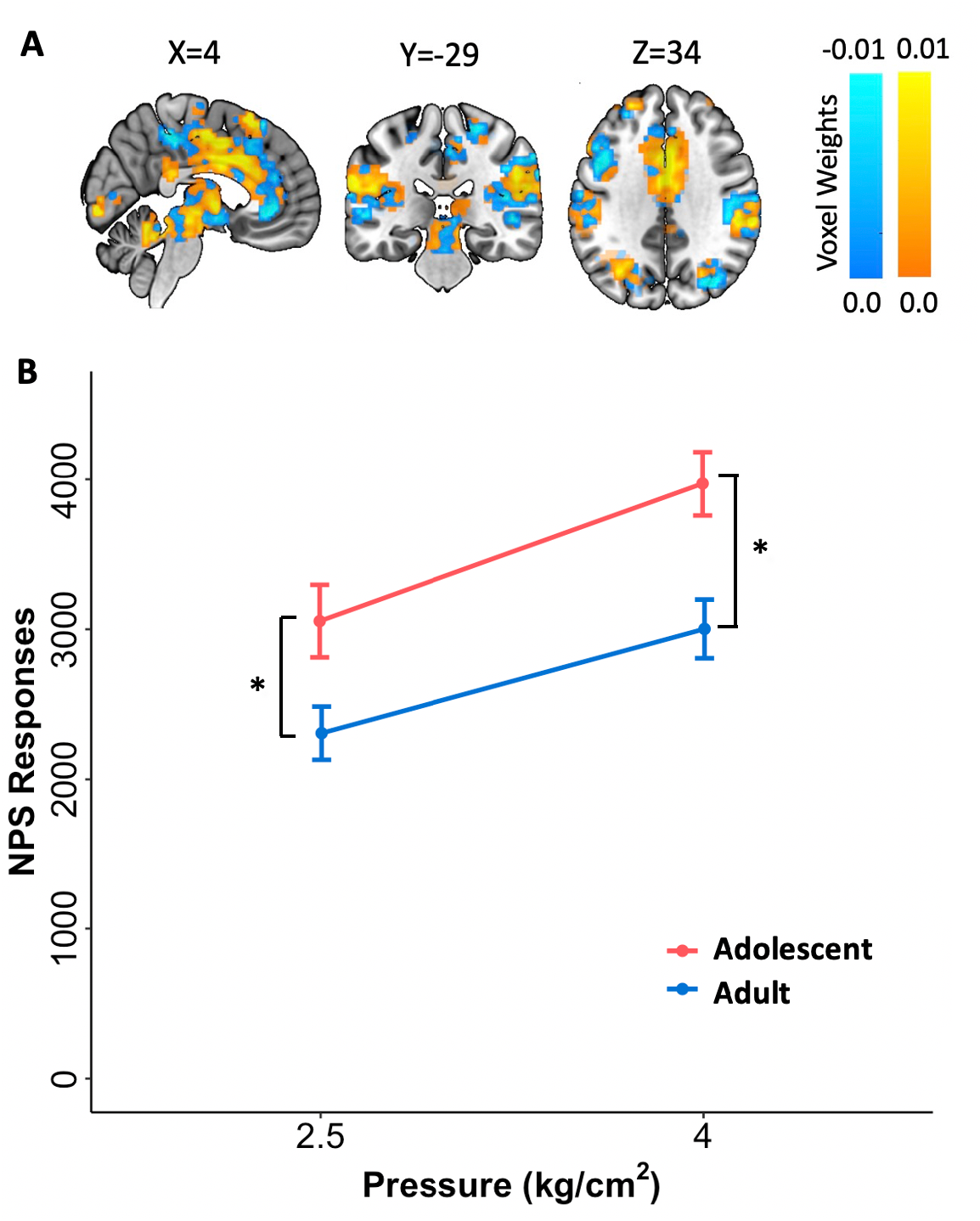
Pain-evoked brain responses in adolescent group, adult group and between-group comparisons. (A) Brain responses to noxious pressure stimuli at 2.5kg/cm2. Adolescents showed greater activation than adults in dorsolateral and dorsomedial PFC, and supramarginal gyrus, along with greater deactivation in the medial orbitofrontal cortex, in response to stimuli at 2.5/cm2. (B) Brain responses to noxious pressure stimuli at 4kg/cm2. Adolescents showed greater activation than adults in rostral anterior cingulate cortex and dorsomedial prefrontal cortex. Clusters of voxels were identified using a threshold of Z>3.1 and their statistical significance (p<0.05) was estimated according to Gaussian random field theory (Worsley KJ et al., 1992). X, Y, Z are MNI coordinates.

#### 3.2.2 Adolescents have stronger Neurologic Pain Signature responses during pain

The NPS is a map of brain voxel weights that is sensitive and specific to physical pain as opposed to other related, yet different, negative experiences[48,56,57,58,94]. As shown in **Figure 4A**, yellow and blue colors were used to represent positive and negative predictive weights respectively. These NPS weights were applied to each participant’s contrast image for the pain period to compute NPS pattern expression. **Figure 4B** shows pain-evoked NPS responses by pressure and group. As expected, the NPS was strongly expressed in both groups during pressure pain at 2.5 kg/cm^2^ (adolescent group: t=13.53, p<0.0001, effect size Cohen’s d=2.47; adult group: t=10.59, p<0.0001, d=1.93) and 4 kg/cm^2^ (adolescent group: t=17.25, p<0.0001, d=3.15; adult group: t=12.37, p<0.0001, d=2.26). Results from the mixed-effects ANOVA showed a significant main effect of group (F=8.04, p=0.006) and pressure (F=48.00, p<0.0001) on NPS responses. Unlike what we found for pain intensity ratings, we did not find an interaction effect (F=0.92, p=0.343) for NPS responses. Post-hoc between-group comparisons showed that adolescents had significantly stronger NPS responses to painful stimuli than adults at both 2.5 kg/cm^2^ (t=2.30, p=0.025, effect size Cohen’s d=0.61) and 4 kg/cm^2^ (t=2.99, p=0.004, d=0.75).

**Figure 4.**
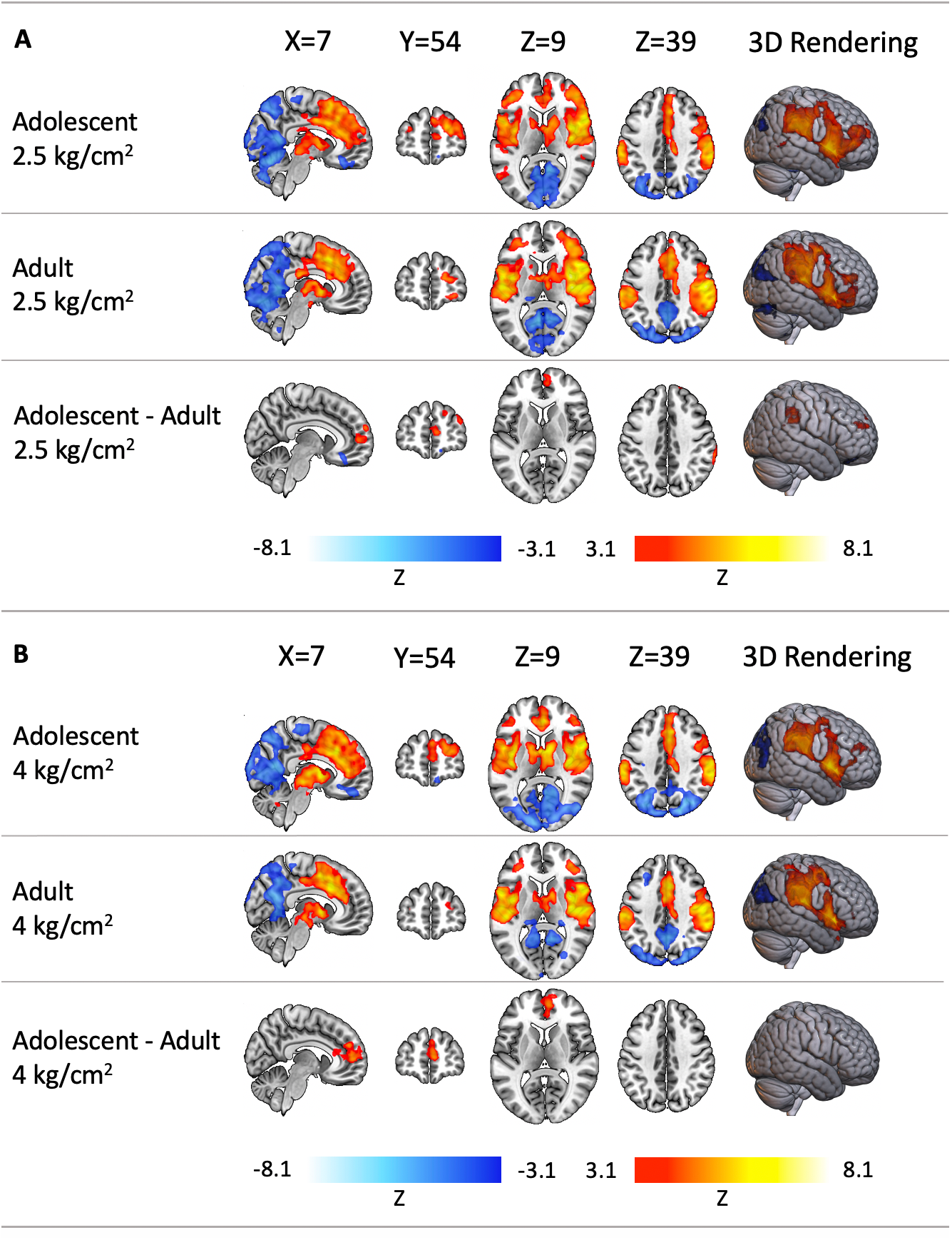
The Neurologic Pain Signature (NPS) pattern and pain-evoked NPS responses. (A) The NPS, an fMRI-based brain signature for physical pain, is a map of brain voxel weights that can predict pain intensity at the individual person level (Wager TD et al., 2013). Voxels in yellow represent positive predictive weights whereas voxels in blue represent negative predictive weights. (B) Both adolescents and adults showed significant pain-evoked NPS responses. Adolescents had greater NPS responses than adults to noxious pressure stimuli at both 2.5kg/cm2 and 4kg/cm2. Error bars represent standard error of the mean. *p<0.05 in post-hoc t-test following mixed-design ANOVA.

#### 3.2.3 Adolescents show augmented pain-evoked neural responses in the default mode network and the ventral attention network

We examined pain-evoked activation differences between groups within seven large-scale cortical resting-state networks as identified in the study by Yeo and colleagues (N=1000 participants)[99]. A single scalar value was computed for each of these seven networks in each participant, respectively, by taking the dot product of contrast images of parameter estimates for the pain period and the binary mask of the network (**Figure 5**). For both pressure pain conditions, significant group activation was found in the somatomotor network, the frontoparietal network, and the ventral attentional network (**Table S7**). In addition, deactivations were found in the dorsal attentional network and the visual network. The default mode network was found to be significantly deactivated only in adults in response to 4 kg/cm^2^ (**Table S7**). Importantly, adolescents showed augmented pain-evoked neural responses in ventral attentional (2.5 kg/cm^2^: t=2.94, p= 0.0048; 4 kg/cm^2^: t=3.07, p=0.0033) and default mode networks (2.5 kg/cm^2^: t=2.14, p=0.0371; 4 kg/cm^2^: t=2.79, p=0.0074) when compared with adults. Adolescents also exhibited greater deactivations in visual network during pain caused by pressure stimuli at 4 kg/cm^2^ (t=2.50, p=0.0155).

**Figure 5.**
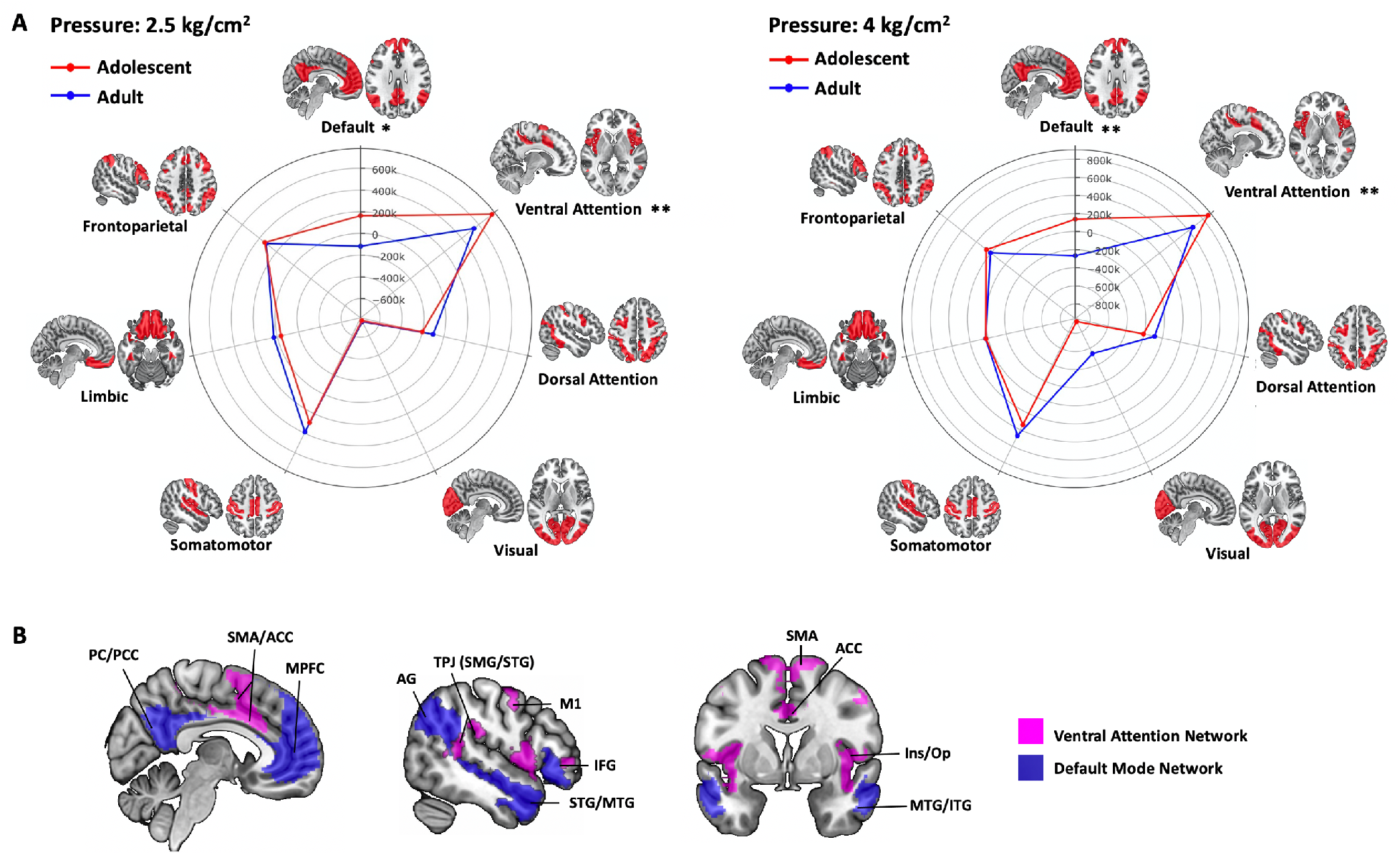
Pain-evoked neural responses within seven major resting-state cortical networks (as described in Yeo BTT et al., 2011) and the brain regions forming the ventral attention network and the default mode network. (A) Polar plots comparing pain-evoked brain responses to noxious pressure stimuli at 2.5 kg/cm2 and 4 kg/cm2 between adolescent group and adult group within 7 major cortical networks. The numerical values are the group means of the dot product of the pre-defined masks of these networks and each participant’s contrast images of parameter estimates for the pain period (2.5 kg/cm2 or 4 kg/cm2). * p<0.05, ** p<0.01 in two-sample t-test. (B) Representation of the brain regions forming ventral attention network and default mode network (Yeo BTT et al., 2011). AC =anterior cingulate cortex, AG=angular gyrus, IFG= inferior frontal gyrus, Ins=insula, ITG= inferior temporal gyrus, MPFC = medial prefrontal cortex, MTG = middle temporal gyrus, M1 = primary motor cortex, Op=Operculum, PC=Precuneus, PCC=posterior cingulate cortex, STG=superior temporal gyrus, SMA=supplementary motor area, SMG=supramarginal gyrus, TPJ=temporoparietal junction.

#### 3.2.4 Pain-evoked brain activation in limbic and prefrontal regions predict and mediate pain perception in adolescents

To identify the brain systems that (1) most strongly predict pain perception in adolescents even after controlling for stimulus intensity and that (2) mediate the effects of noxious stimulus intensity on pain perception in adolescents, we conducted whole-brain multilevel mediation analyses across trial-by-trial estimates of brain and behavioral responses during pain[4,5,45,56]. **Figure 6** shows a diagram of the mediation model.

**Figure 6.**
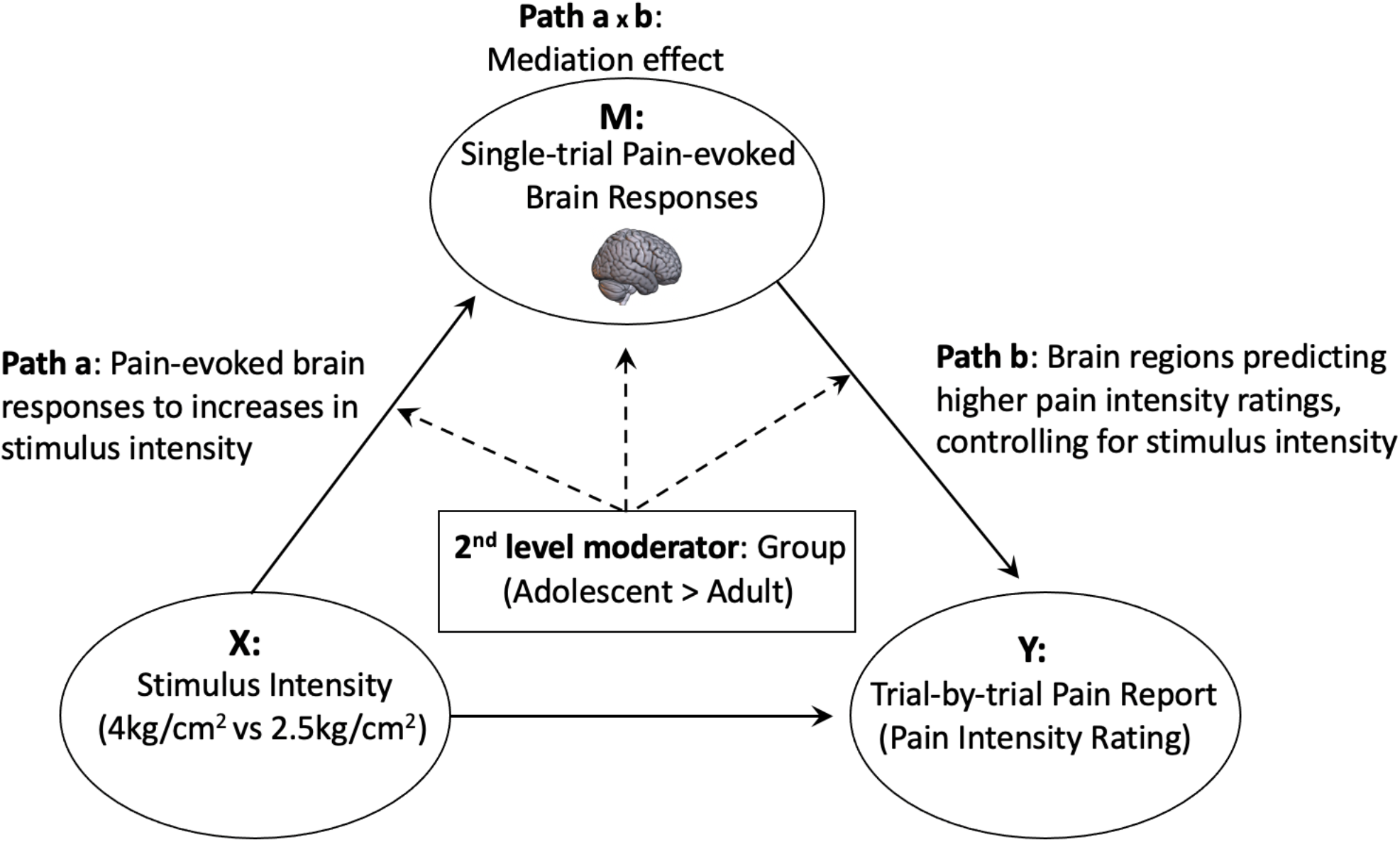
Whole-brain multilevel mediation model, with stimulus intensity as the predictor, single trial pain-evoked brain activity as the mediating factor, and pain intensity ratings as the outcome. Group (adolescent vs. adult) was included as the second-level moderator to investigate adolescence induced changes.

The results for path b in adolescents showed that greater activation of the amygdala and parahippocampal gyrus bilaterally significantly predicted greater pain perception above and beyond the effects of stimulus intensity (**Figure 7A**). Other significant regions for path b in adolescents included the posterior insula, secondary somatosensory cortex, primary sensorimotor cortex in the paracentral lobule, dorsolateral prefrontal cortex, midcingulate cortex, temporal cortex, lateral occipital cortex and putamen (**Table S8**). Interestingly, we did not find pain-evoked neural responses in amygdala and parahippocampal gyrus as strong predictors of greater pain perception (path b effect) in adults (**Figure 7B** and **Table S9**). Furthermore, results from the second-level moderator analysis showed that the bilateral parahippocampal gyrus and clusters in the amygdala/hippocampus, midcingulate cortex, paracentral lobule, premotor cortex and temporal cortex were significantly stronger predictors of pain intensity in adolescents than in adults (**Figure 7C** and **Table S10**). For the second level moderator analyses, we chose a more lenient uncorrected p<0.001 threshold at the voxel level as in previous studies[45].

**Figure 7.**
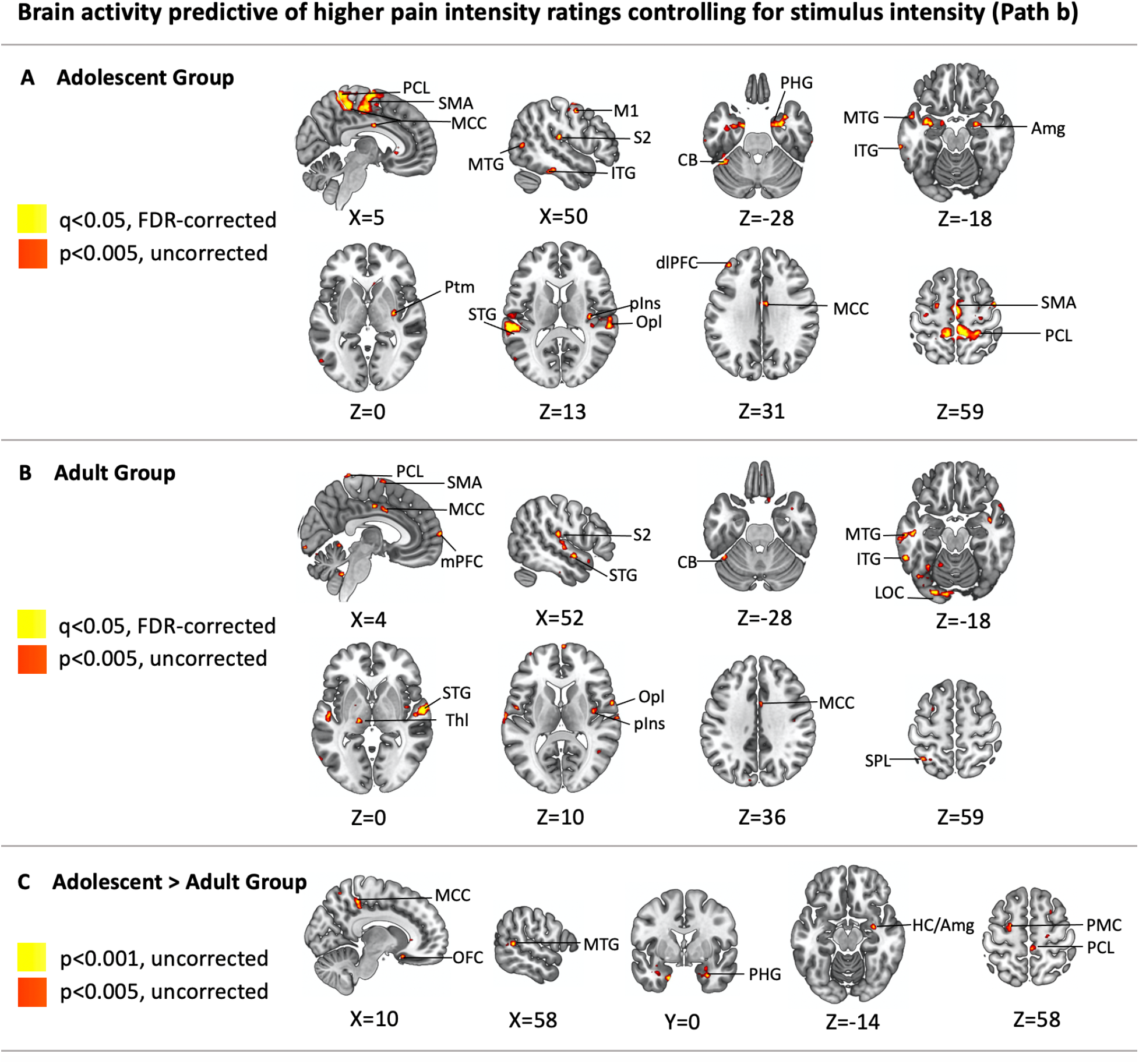
Brain activity predictive of higher pain intensity ratings controlling for stimulus intensity. (A) Brain predictors for pain intensity ratings in adolescents (Path b effect). (B) Brain predictors for pain intensity ratings in adults (Path b effect). (C) Differences between adolescents and adults in brain predictors of higher pain intensity ratings (group moderated Path b effect: Adolescent > Adult). PCL=paracentral lobule, SMA=supplemental motor area, M1=primary motor cortex, S2=secondary somatosensory cortex, ITG=inferior temporal gyrus, CB=cerebellum, PHG=parahippocampus, Amg=amygdala, Ptm=putamen, STG=superior temporal gyrus, pIns=posterior insula, Opl=operculum, dlPFC=dorsolateral prefrontal cortex, MCC= medial cingulate cortex, mPFC=medial prefrontal cortex, MTG=middle temporal gyrus, LOC=lateral occipital cortex, Thl=thalamus, SPL=superior parietal lobule, OFC=orbitofrontal cortex, HC=hippocampus, PMC=premotor cortex.

The results for path a x b in adolescents showed that the brain mediators of noxious stimulus intensity on pain perception involved mostly regions that were significantly activated during pain, including the amygdala/hippocampus, parahippocampal gyrus, prefrontal regions, midcingulate cortex, supramarginal gyrus, and ventral striatum (**Figure 8A** and **Table S11**). The observed mediation effect in these regions indicates that greater increases in pain-evoked activation during high vs. low pressure in such regions were also predictive of larger increases in pain intensity ratings (even after controlling for pressure intensity) in adolescents. The results for path a x b in adults seem a bit more spatially scattered when visually compared with adolescents, but does not include dorsomedial prefrontal cortex and parahippocampal gyrus (**Figure 8B** and **Table S12**). Importantly, clusters within the dorsomedial PFC and right ventrolateral PFC, parahippocampal gyrus, midcingulate cortex and temporal cortex showed a significant moderator effect (**Figure 8C** and **Table S13**), indicating that these regions were stronger mediators of subjective pain perception in adolescents than in adults. Consistent with our previous GLM results showing augmented activation of the medial and lateral PFC in adolescents than in adults, these findings suggest a role for these regions in more strongly contributing to pain perception in adolescents.

**Figure 8.**
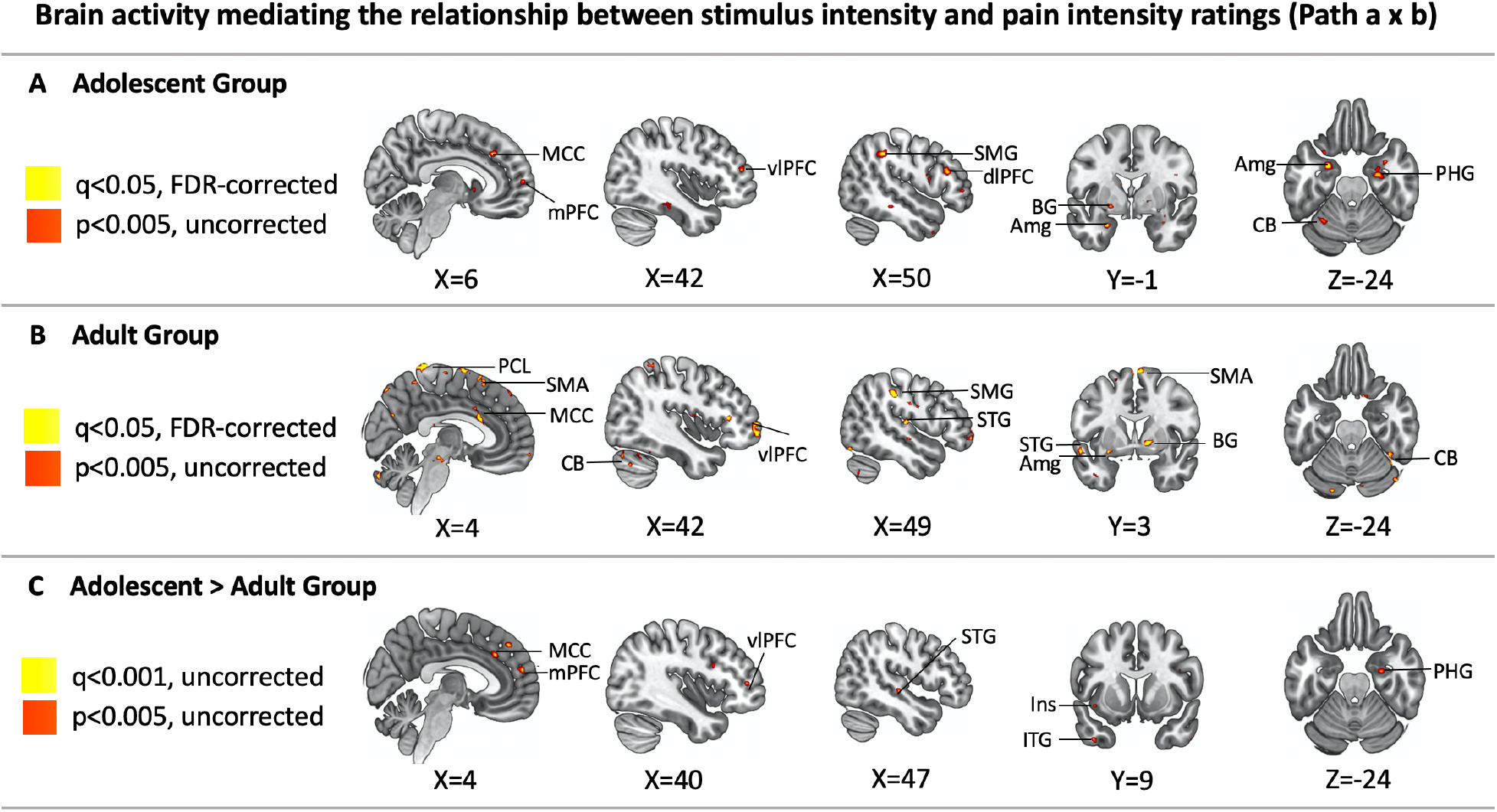
Brain activity mediating the relationship between stimulus intensity and pain intensity ratings. (A) Brain mediators of higher pain intensity ratings to stimuli with greater intensity in adolescents (Path a x b effect). (B) Brain mediators of higher pain intensity ratings to stimuli with greater intensity in adults (Path ab effect). (C) Differences between adolescents and adults in brain activity mediating the relationship between stimulus intensity and pain intensity ratings (group moderated Path a x b effect: Adolescent > Adult). MCC=medial cingulate cortex, mPFC=medial prefrontal cortex, vlPFC=ventrolateral prefrontal cortex, SMG=supramarginal gyrus, dlPFC=dorsolateral prefrontal cortex, PHG=parahippocampus, BG=basal ganglia, Amg=amygdala, CB=cerebellum, SMA= supplemental motor area, STG=superior temporal gyrus, Ins= insula, ITG=inferior temporal gyrus.

## Discussion

To our knowledge, this is the first study that directly compares pain perception and brain responses to acute experimental noxious stimuli between adolescents and adults. We found that, compared to adult women, adolescent females were more sensitive to painful pressure at low stimulus intensities and showed remarkably stronger pain-related responses of NPS, an fMRI-based brain marker for acute physical pain perception[94]. We also found that regions within the medial prefrontal cortex, the default mode network, the amygdala and associated hippocampal and striatal regions were more strongly activated during pain or showed a greater contribution to predicting pain experience in adolescents. Taken together, the findings suggest that adolescence particularly in females is a developmental period characterized by increased sensitivity to pain, potentially through two mechanisms: (1) augmented nociceptive signal processing at the central nervous system (CNS) level, which may reflect (at least in part) augmented peripheral input to the CNS and (2) augmented involvement of core brain regions for aversive emotion appraisal, regulation, affective learning and memory. The hyper-representation of acute pain in the adolescent female brain may underlie augmented vulnerability to acute painful experiences and associated aversive memories during adolescence. Futures studies are warranted to further establish this association, its underlying neurobiology and its relationship with the steep increase of bodily pains that is observed, particularly in females, in the transition to adolescence.

We found a group by stimulus intensity interaction effect predicting pain intensity ratings, suggesting that the heightened pain sensitivity in adolescents is stimulus-intensity-dependent. Specifically, adolescents reported greater pain intensity and unpleasantness than middle-aged adults in response to low-intensity peri-threshold noxious stimuli (at 2.5 kg/cm^2^). This finding is in line with the observation that pain threshold generally increases with age[92]. It suggests that adolescents are more sensitive to noxious pressure than adults at low stimulus intensities. However, we also observed that this difference in pain perception between adolescents and adults disappeared as the stimulus intensity increased to 4 kg/cm^2^. The underlying mechanisms for increased sensitivity to low noxious pressure in adolescents could be related to a greater density of nociceptor-containing sensory nerve fibers found in their skin or deep tissue[42, 53]. However, this possibility does not readily explain the observed stimulus intensity dependence of pain sensitivity in adolescents. The mechanisms might also involve the central nervous system, specifically the brain, where the pain perception is generated and modulated.

The standard massive univariate GLM analyses showed that adolescents exhibited greater pain-evoked activation in the PFC (medial and middle frontal gyrus) and the PPC (supramarginal gyrus) in response to low-intensity noxious pressure. The PFC and the PPC are often activated during acute experimental pain[3,21,46,68,91], and have been associated with cognitive aspects of pain perception such as spatial attention and evaluation of the spatial location of noxious stimuli[54, 69]. Both regions are part of the association cortex that is undergoing dynamic maturation during adolescence through synaptic pruning[12,31,47]. Our finding is consistent with the results of previous fMRI studies showing greater PFC and PPC activation in adolescents than in adults during cognitive tasks[13, 61]. The increased pain-evoked brain activation of these brain regions might be associated with the firing of an excessive number of synapses that are still waiting to be pruned. It may also reflect, at least in part, a compensatory brain response to more nociceptive input.

We then compared the pain-evoked responses in the NPS, an fMRI-based spatial and magnitude pattern for perception of acute physical pain[94], between adolescents and adults. Adolescents showed stronger NPS responses to both low and high levels of noxious pressure than adults (i.e., 2.5 kg/cm^2^ and 4 kg/cm^2^). We interpret this finding as suggesting that adolescents have an overall increase in nociception-related signal processing in the brain. Again, the underlying mechanisms may involve adolescents’ relative hypersensitivity in the central and/or peripheral nervous system. Interestingly, we did not find a group by stimulus intensity interaction effect for NPS responses as we found for subjective pain ratings. This implies that the augmented sensitivity to lower stimulus intensities in adolescents may involve pain-related neural processes not reflected in NPS.

To further identify these processes, we compared pain-evoked neural responses within each of the seven previously identified large-scale cortical networks[99]. We found that adolescents showed augmented responses within the DMN and the ventral attention network (VAN). The DMN is composed of medial PFC, the posterior cingulate cortex (PCC)/precuneus, the lateral parietal cortex, and parahippocampal gyrus and characterized by being active when a person is at rest and being deactivated during externally-oriented tasks[8,28,33,79,85]. Regions of DMN, particularly the medial PFC, are also found to be activated during internal mentation such as autobiographical memory recall[11,62,89] and tasks associated with social or self-referential processing[29, 81]. Core regions of DMN (medial PFC, PCC) are typically deactivated during acute experimental pain[2, 46]. The paradoxical pain-evoked activation of medial PFC in adolescents could reflect augmented self-referential processing while they experience pain, possibly associated with episodes of recollection of past or formation of new memories of physical pain. The VAN includes regions in the right-lateralized temporo-parietal junction (including supramarginal gyrus and superior temporal gyrus), ventrolateral frontal cortex, anterior insula and anterior cingulate cortex, and is typically activated by salient sensory stimuli, such as pain[18,19,67]. This network is also often known as the salience network[49, 82]. The VAN has been functionally associated with breaking one’s attention from the current task and reorienting it to an unexpected salient external stimuli (i.e. bottom-up processing)[18, 84]. The observed increased pain-evoked VAN responses in adolescents may suggest greater attentional demand during pain response than in adults. This might be interpreted as immature, less-efficient functioning of the associative cortices that encompass the VAN. This could also suggest a physiological response to augmented nociceptive input.

Lastly, using the statistically robust multilevel mediation approach[4,5,45,56], we explored the relationships between stimulus intensity, single-trial pain-evoked brain responses, and single-trial pain ratings. We focused on brain predictors of pain experience controlling for stimulus intensity (Path b) and brain activity mediating the relationship between pressure intensity and subjective pain experience (Path a x b), since those are the two paths in the model directly linking brain responses to subjective experience. Our results showed that activations in the amygdala and associated regions (i.e., hippocampus, parahippocampal gyrus), which are pivotal for emotional processing and formation of aversive memories[72, 80], played a stronger role in adolescents compared with adults in predicting higher pain intensity ratings. In addition, we found that adolescents’ increased activity in key regions comprising DMN (medial PFC, parahippocampus, inferior temporal cortex) and VAN (ventrolateral PFC, insula, anterior cingulate, supramarginal gyrus, superior temporal gyrus) mediated the between-group difference in the relationship between stimulus intensity and pain intensity ratings. Overall, these findings complement and reinforce previous findings showing that limbic regions, together with regions that are involved in self-referential processing and bottom-up attentional reorienting, mediate subjective pain experience in adolescents.

In conclusion, this study provides the first evidence of augmented pain-evoked brain responses in healthy adolescent females involving regions important for affective, cognitive as well as nociceptive processing, which is compatible with their heightened pain sensitivity to low-intensity noxious stimuli, compared to adult women. The present results also confirm that age represents a significant source of individual differences in perceived pain as well as noxious stimulus-related brain activation. Further studies are needed to examine sex differences in brain responses to pain and ages at which these differences begin to unfold. The observation that different patterns of activations were noted between adolescents and adults has important implications for the development of neuroimaging-based markers for pain intensity. Will different markers of pain intensity be needed to accurately assess inter-individual differences in pain across different age groups? If such developmentally specific markers are required, would separate markers be required for infants, prepubertal children, adolescents, adults, as well as elderly individuals? What implications do these findings have for assessment of pain in clinical populations? A greater emphasis of developmentally-informed research on pain across the entire lifespan is clearly needed.

## Acknowledgements

The authors have no conflict of interest to declare.

This work was funded by Cincinnati Children’s Hospital Medical Center’s Trustee Grant Award and NIH/NIAMS Grants R01 AR074795 and P30 AR076316. Marina López-Solà, Ph.D. is hired as part of the Serra Hunter Programme of the Generalitat de Catalunya.

The authors gratefully thank Matt Lanier, Kaley Bridgewater, Kelsey Murphy, Brynne Williams, and Lacey Haas (Imaging Research Center, Department of Radiology, Cincinnati Children’s Hospital Medical Center) for their contributions to MRI data collection.

