## Supplemental Materials for "Processing of Pain by the Developing Brain: Evidence of Differences Between Adolescent and Adult Females"

**Supplemental Figures**

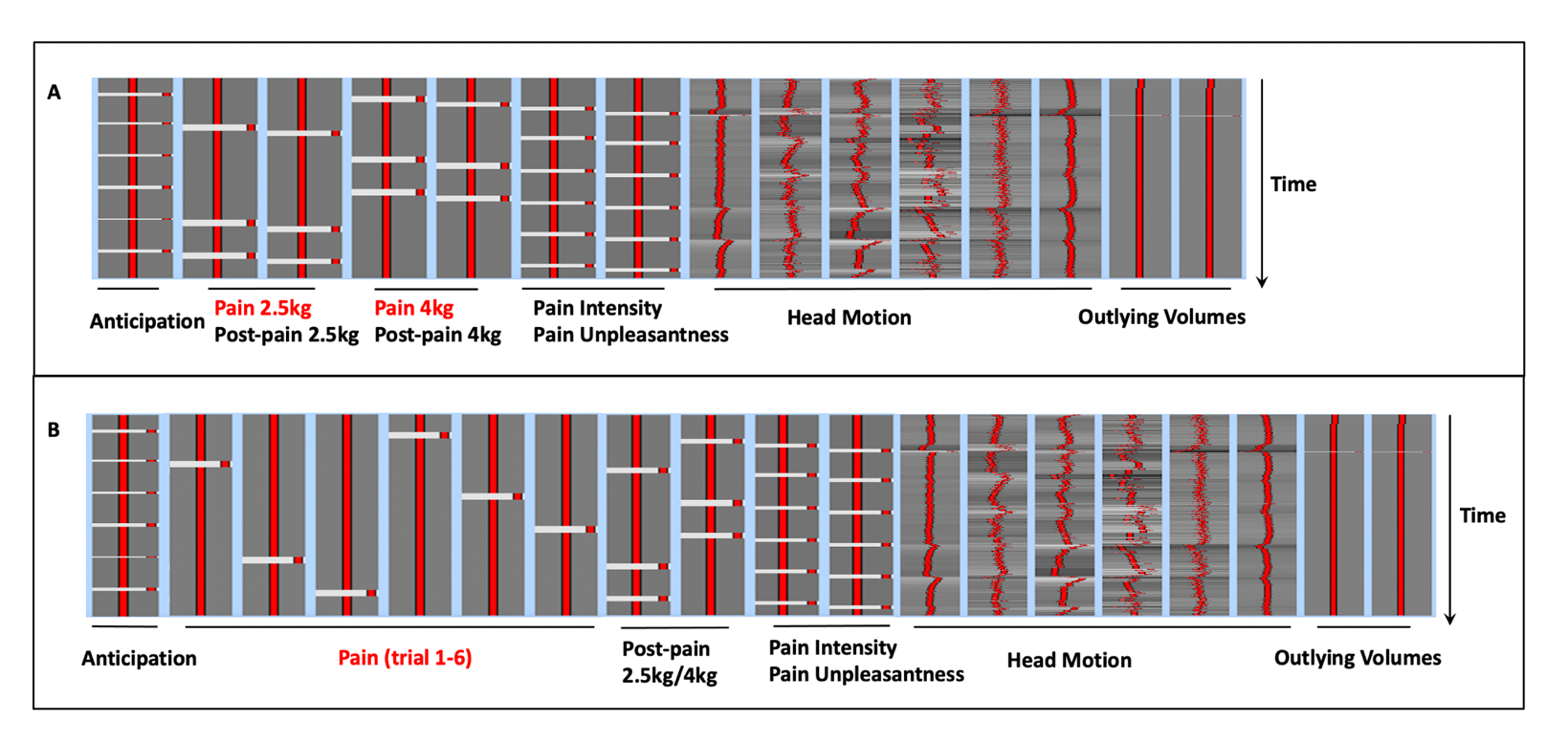

**Figure S1** Two ways of study design implemented in the first-level general linear model analyses. **(A)** The traditional model included 2 pain-period regressors, one for 3 trials associated with noxious stimuli at 2.5 kg/cm^2^, and one for the 3 trials associated with stimuli at 4 kg/cm^2^. **(B)** The trial-by-trial model included 6 pain period regressors for all 6 trials in one fMRI run. The results were used in multilevel mediation analyses. Regressors for pain periods were highlighted in red.

**Supplemental Tables**

**Table S1. Brain responses to 2.5kg/cm2 stimuli in the adolescent group (Z>3.1, p<0.05, cluster-corrected).**

| **Activation** |  |  |  |  |  |
| --- | --- | --- | --- | --- | --- |
| **Cluster 1** | **Voxel Size** | 36324 | **P-value** | 2.33E-24 |  |
| **Region** | **Side** | **X** | **Y** | **Z** | **Z-Score** |
| Central operculum | R | 54 | 2 | 2 | 8.16 |
| Central operculum | L | -54 | -2 | 4 | 5.98 |
| Temporal Pole | R | 52 | 12 | -2 | 7.92 |
| Temporal Pole | L | -42 | 4 | -16 | 6.58 |
| Middle insula | R | 40 | -4 | -8 | 7.41 |
| Middle insula | L | -40 | 2 | -4 | 6.95 |
| Anterior insula | R | 38 | 16 | 0 | 6.95 |
| Anterior insula | L | -42 | 12 | -6 | 6.97 |
| Posterior insula | R | 40 | -12 | 4 | 6.41 |
| Posterior insula | L | -38 | -16 | 0 | 5.8 |
| Parietal operculum (S2) | R | 54 | -26 | 18 | 7.18 |
| Parietal operculum (S2) | L | -54 | -26 | 20 | 6.67 |
| Supramarginal gyrus | L | -64 | -26 | 24 | 7.15 |
| Supramarginal gyrus | R | 62 | -24 | 24 | 7.09 |
| Amygdala | R | 24 | 2 | -14 | 6.18 |
| Amygdala | L | -18 | 4 | -16 | 7.04 |
| Postcentral gyrus (S1) | R | 66 | -18 | 30 | 6.46 |
| Postcentral gyrus (S1) | L | -64 | -14 | 20 | 5.95 |
| Inferior frontal gyrus (BA46) | R | 48 | 40 | 0 | 6.4 |
| Inferior frontal gyrus (BA46) | L | -40 | 40 | 14 | 5.73 |
| Precentral gyrus (M1) | R | 56 | 8 | 22 | 6.23 |
| Precentral gyrus (M1) | L | -54 | 6 | 4 | 6.11 |
| Supplemental Motor Area (BA6) | R | 12 | 12 | 62 | 5.97 |
| Anterior Cingulate Cortex | R | 6 | 14 | 32 | 5.85 |
| Anterior Cingulate Cortex | L | -8 | 36 | 14 | 5.55 |
| Superior Frontal Gyrus (BA10) | R | 22 | 58 | 20 | 5.78 |
| Middle frontal Gyrus (BA10) | L | -38 | 44 | 22 | 5.69 |
| Putamen | R | 28 | 12 | -4 | 5.7 |
| Caudate | R | 12 | 6 | 10 | 5.44 |
| Caudate | L | -15 | 2 | 15 | 5.01 |
| Medial Superior Frontal Gyrus (BA8) | R | 6 | 32 | 50 | 5.3 |
| Posterior Cingulate Cortex | R | 12 | -26 | 38 | 5.11 |
| Periaqueductal gray | N/A | 4 | -24 | -4 | 4.97 |
| Middle temporal gyrus | R | -50 | -60 | 4 | 4.83 |
| Middle temporal gyrus | R | 58 | -22 | -10 | 4.75 |
| Thalamus | R | 6 | -18 | 2 | 4.3 |
| **Deactivation** |  |  |  |  |  |
| **Cluster 1** | **Voxel Size** | 20285 | **P-value** | 1.15E-16 |  |
| **Region** | **Side** | **X** | **Y** | **Z** | **Z-Score** |
| Fusiform Gyrus | R | 32 | -42 | -24 | 6.21 |
| Fusiform Gyrus | L | -36 | -54 | -16 | 6.14 |
| Cerebellum | R | 14 | -54 | -20 | 6.18 |
| Cerebellum | L | -20 | -48 | -20 | 4.82 |
| Cerebellum Vermis | N/A | 4 | -66 | -20 | 6.13 |
| Precuneus | R | 8 | -54 | 10 | 5.89 |
| Precuneus | L | -10 | -60 | 12 | 4.83 |
| Lateral occipital cortex | L | -36 | -76 | 32 | 5.39 |
| Lateral occipital cortex | R | 44 | -78 | 30 | 5.22 |
| Precentral Gyrus (M1) | L | -42 | -18 | 64 | 5.38 |
| Postcentral Gyrus (S1) | L | -8 | -38 | 66 | 5.32 |
| Pons | L | -2 | -14 | -36 | 5.12 |
| Intracalcarine cortex | L | -1 | -84 | 6 | 4.82 |
| Intracalcarine cortex | R | 12 | -84 | 6 | 4.2 |
| **Cluster 2** | **Voxel Size** | 665 | **P-value** | 0.038 |  |
| **Region** | **Side** | **X** | **Y** | **Z** | **Z-Score** |
| Rectus gyrus (BA11) | L | -8 | 28 | -18 | 4.82 |
| Rectus gyrus (BA11) | R | 10 | 30 | -18 | 4.44 |

**Table S2. Brain responses to 2.5kg/cm2 stimuli in the adult group (Z>3.1, p<0.05, cluster-corrected).**

| **Activation** |  |  |  |  |  |
| --- | --- | --- | --- | --- | --- |
| **Cluster 1** | **Voxel Size** | 41101 | **P-value** | 1.18E-25 |  |
| **Region** | **Side** | **X** | **Y** | **Z** | **Z-Score** |
| Middle insula | R | 42 | 0 | 0 | 7.82 |
| Middle insula | L | -42 | -2 | 8 | 6.38 |
| Anterior insula | R | 38 | 20 | -6 | 7.58 |
| Anterior insula | L | -32 | 14 | 0 | 6.89 |
| Parietal operculum | R | 54 | -26 | 18 | 7.38 |
| Parietal operculum | L | -50 | -32 | 16 | 5.69 |
| Superior temporal gyrus | R | 54 | 4 | -2 | 7.15 |
| Superior temporal gyrus | L | -48 | -26 | 10 | 6.55 |
| Posterior insula | R | 40 | -16 | 12 | 7.13 |
| Posterior insula | L | -40 | -18 | 10 | 5.1 |
| Central Operculum | R | 60 | -14 | 14 | 7.11 |
| Central Operculum | L | -58 | -18 | 14 | 6.71 |
| Postcentral gyrus | R | 54 | -16 | 38 | 6.96 |
| Postcentral gyrus | L | -58 | -18 | 28 | 6.76 |
| Amygdala | R | 22 | 4 | -16 | 6.8 |
| Amygdala | L | -20 | 2 | -18 | 5.81 |
| Inferior frontal gyrus | R | 48 | 16 | 8 | 6.79 |
| Supramarginal gyrus | R | 60 | -22 | 32 | 6.71 |
| Supramarginal gyrus | L | -66 | -26 | 38 | 5.86 |
| Orbital frontal cortex | R | 42 | 38 | -10 | 6.62 |
| Orbital frontal cortex | L | -30 | 18 | -16 | 5.43 |
| Precentral gyrus | R | 46 | -2 | 54 | 6.43 |
| Precentral gyrus | L | -54 | 4 | 6 | 5.96 |
| Anterior cingulate cortex | R | 6 | 12 | 44 | 6.19 |
| Anterior cingulate cortex | L | -1 | 8 | 30 | 5.06 |
| Putamen | R | 20 | 10 | -8 | 6.11 |
| Putamen | L | -18 | 4 | -8 | 6.03 |
| Superior frontal gyrus (BA6) | R | 14 | 12 | 60 | 5.86 |
| Superior frontal gyrus (BA6) | L | -14 | 22 | 56 | 3.99 |
| Superior frontal gyrus (BA10) | R | 20 | 50 | 18 | 5.79 |
| Middle frontal gyrus (BA10) | R | 38 | 42 | 10 | 5.75 |
| Middle frontal gyrus (BA10) | L | -38 | 42 | 18 | 4.69 |
| Caudate | R | 14 | 16 | -4 | 5.71 |
| Caudate | L | -10 | 18 | -2 | 5.46 |
| Thalamus | R | 6 | -4 | 12 | 5.48 |
| Thalamus | L | -4 | -4 | 10 | 5.31 |
| Supplementary motor area | R | 6 | 0 | 48 | 5.43 |
| Supplementary motor area | L | -6 | 6 | 50 | 5.31 |
| Posterior cingulate cortex | R | 12 | -20 | 36 | 4.88 |
| Hippocampus | R | 20 | -14 | -18 | 4.13 |
| Periaqueductal gray | N/A | 2 | -28 | -6 | 3.7 |
| **Cluster 2** | **Voxel Size** | 1464 | **P-value** | 0.00334 |  |
| Cerebellum | L | -22 | -72 | -50 | 6.01 |
| **Deactivation** |  |  |  |  |  |
| **Cluster 1** | **Voxel Size** | 24741 | **P-value** | 2.00E-18 |  |
| **Region** | **Side** | **X** | **Y** | **Z** | **Z-Score** |
| Fusiform Gyrus | R | 34 | -38 | -22 | 6.85 |
| Fusiform Gyrus | L | -42 | -50 | -24 | 5.47 |
| Lingual gyrus | R | 10 | -54 | 0 | 5.78 |
| Lingual gyrus | L | -14 | -44 | -6 | 4.41 |
| Precuneus | R | 2 | -56 | 46 | 5.17 |
| Precuneus | L | 0 | -70 | 28 | 5.58 |
| Cerebellum | R | 6 | -72 | -24 | 5.11 |
| Postcentral gyrus | R | 4 | -42 | 64 | 4.03 |
| Occipital Pole | R | 8 | -94 | -10 | 4.84 |
| Occipital Pole | L | -6 | -98 | 4 | 4.2 |

**Table S3. Brain responses to 4kg/cm2 stimuli in the adolescent group (Z>3.1, p<0.05, cluster-corrected).**

| **Activation** |  |  |  |  |  |
| --- | --- | --- | --- | --- | --- |
| **Cluster 1** | **Voxel Size** | 35360 | **P-value** | 1.57E-24 |  |
| **Region** | **Side** | **X** | **Y** | **Z** | **Z-Score** |
| Central operculum | R | 54 | 4 | 0 | 8.11 |
| Middle insula | R | 42 | 8 | -4 | 8.1 |
| Middle insula | L | -42 | 2 | -6 | 7.92 |
| Anterior insula | L | -44 | 16 | -6 | 7.76 |
| Anterior insula | R | 36 | 24 | -2 | 7.45 |
| Superior temporal gyrus | L | -42 | -10 | -12 | 7.71 |
| Superior temporal gyrus | R | 42 | -2 | -20 | 7.34 |
| Supramarginal gyrus | R | 62 | -24 | 24 | 7.62 |
| Supramarginal gyrus | L | -62 | -26 | 20 | 7.26 |
| Posterior insula | R | 40 | -10 | -6 | 7.45 |
| Posterior insula | L | -38 | -18 | 10 | 6.28 |
| Parietal Operculum | R | 54 | -26 | 20 | 6.98 |
| Amygdala | R | 26 | 4 | -16 | 6.9 |
| Amygdala | L | -22 | 4 | -14 | 6.22 |
| Postcentral gyrus | L | -60 | -18 | 20 | 6.74 |
| Postcentral gyrus | R | 66 | -18 | 32 | 6.23 |
| Anterior Midcingulate cortex | R | 6 | 24 | 36 | 6.36 |
| Anterior Midcingulate cortex | L | -4 | 24 | 28 | 5.2 |
| Pregenual anterior cingulate cortex | R | 4 | 40 | 8 | 6.03 |
| Pregenual Anterior cingulate cortex | L | -4 | 36 | 18 | 5.64 |
| Medial superior frontal gyrus | R | 4 | 36 | 46 | 6.11 |
| Thalamus | R | 14 | -4 | 4 | 6 |
| Thalamus | L | -8 | -12 | 4 | 5.1 |
| Pallidum/Putamen | R | 16 | 8 | -2 | 5.89 |
| Periaqueductal gray | N/A | 6 | -22 | -6 | 5.77 |
| Supplementary motor cortex | R | 10 | 24 | 60 | 5.65 |
| Inferior frontal gyrus | R | 46 | 38 | 0 | 5.71 |
| Inferior frontal gyrus | L | -38 | 40 | 12 | 4.41 |
| Mddile frontal gyrus | R | 40 | 44 | 24 | 5.09 |
| Middle frontal gyrus | L | -38 | 40 | 24 | 5.62 |
| Accumbens/Caudate | R | -6 | 6 | -4 | 5.04 |
| Pons | N/A | 4 | -18 | -22 | 4.28 |
| **Cluster 2** | **Voxel Size** | 1542 | **P-value** | 0.00199 |  |
| **Region** | **Side** | **X** | **Y** | **Z** | **Z-Score** |
| Cerebellum | L | -20 | -70 | -54 | 5.62 |
| Cerebellum Vermis | N/A | 0 | -56 | -38 | 5.48 |
| Cerebellum | R | 14 | -58 | -36 | 4.9 |
| **Deactivation** |  |  |  |  |  |
| **Cluster 1** | **Voxel Size** | 25415 | **P-value** | 9.04E-20 |  |
| **Region** | **Side** | **X** | **Y** | **Z** | **Z-Score** |
| Fusiform gyrus | L | -32 | -58 | -14 | 7.27 |
| Fusiform gyrus | R | 34 | -42 | -26 | 6.34 |
| Lateral occipital cortex | R | 24 | -78 | 44 | 6.48 |
| Lateral occipital cortex | L | -32 | -76 | 18 | 6.33 |
| Precuneous | R | 22 | -56 | 12 | 6.01 |
| Precuneous | L | -10 | -62 | 14 | 4.26 |
| Cerebellum | R | 12 | -58 | -12 | 5.94 |
| Lingual gyrus | R | 22 | -56 | -6 | 5.93 |
| Lingual gyrus | L | -30 | -52 | -6 | 5.89 |
| Occipital Pole | L | -10 | -94 | 8 | 5.64 |
| Occipital Pole | R | 8 | -98 | 20 | 5.56 |
| Intracalcarine cortex | R | 8 | -82 | 4 | 5.13 |
| Intracalcarine cortex | L | -10 | -62 | 8 | 4.09 |
| **Cluster 2** | **Voxel Size** | 1868 | **P-value** | 0.000781 |  |
| **Region** | **Side** | **X** | **Y** | **Z** | **Z-Score** |
| Precentral gyrus | L | -38 | -22 | 70 | 5.74 |
| Postcentral gyrus | L | -40 | -22 | 56 | 5.58 |
| **Cluster 3** | **Voxel Size** | 949 | **P-value** | 0.0132 |  |
| **Region** | **Side** | **X** | **Y** | **Z** | **Z-Score** |
| Rectus gyrus | L | -6 | 30 | -18 | 4.76 |
| Rectus gyrus | R | 8 | 48 | -22 | 4.5 |
| **Cluster 4** | **Voxel Size** | 748 | **P-value** | 0.0273 |  |
| **Region** | **Side** | **X** | **Y** | **Z** | **Z-Score** |
| Paracentral Lobule (M1/S1) | R | 6 | -26 | 64 | 4.58 |
| Paracentral Lobule (M1/S1) | L | -8 | -34 | 68 | 4.47 |

**Table S4. Brain responses to 4kg/cm^2^ stimuli in the adult group (Z>3.1, p<0.05, cluster-corrected).**

| **Activation** |  |  |  |  |  |
| --- | --- | --- | --- | --- | --- |
| **Cluster 1** | **Voxel Size** | 28715 | **P-value** | 1.17E-24 |  |
| **Region** | **Side** | **X** | **Y** | **Z** | **Z-Score** |
| Anterior insula | R | 42 | 16 | -10 | 7.73 |
| Anterior insula | L | -40 | 14 | -8 | 7.24 |
| Central operculum | L | -58 | -18 | 14 | 7.48 |
| Central operculum | R | 58 | -14 | 12 | 7.02 |
| Parietal operculum | R | 52 | -26 | 16 | 7.48 |
| Middle insula | R | 38 | 4 | 6 | 7.47 |
| Middle insula | L | -40 | 0 | -12 | 7.13 |
| Superior temporal gyrus | L | -40 | -10 | -12 | 7.47 |
| Superior temporal gyrus | R | 40 | -6 | -14 | 7.29 |
| Posterior insula | R | 40 | -16 | 12 | 7.23 |
| Posterior insula | L | -44 | -18 | 8 | 6.21 |
| Supramarginal gyrus | R | 64 | -28 | 32 | 7.21 |
| Supramarginal gyrus | L | -64 | -26 | 20 | 6.69 |
| Postcentral gyrus | L | -58 | -18 | 26 | 6.86 |
| Postcentral gyrus | R | 56 | -14 | 44 | 6.79 |
| Orbitofrontal cortex | R | 42 | 28 | -4 | 6.84 |
| Orbitofrontal cortex | L | -44 | 18 | -8 | 6.63 |
| Precentral gyrus | L | -56 | 4 | 6 | 6.54 |
| Precentral gyrus | R | 56 | 10 | 8 | 6.51 |
| Amygdala | R | 22 | 2 | -16 | 6.49 |
| Amygdala | L | -20 | 2 | -20 | 5.46 |
| Putamen | L | -24 | 10 | -8 | 6.08 |
| Putamen | R | 24 | 16 | -4 | 6.05 |
| Middel frontal gyrus | R | 38 | 42 | 12 | 5.25 |
| Middel frontal gyrus | L | -32 | 42 | 16 | 5.02 |
| Periaqueductal gray | N/A | 4 | -20 | -4 | 5.12 |
| Thalamus | R | 14 | -12 | 0 | 4.81 |
| Thalamus | L | -2 | -8 | 12 | 4.3 |
| Caudate | R | 10 | 10 | 0 | 5.06 |
| Caudate | L | -10 | 6 | 6 | 4.52 |
| **Cluster 2** | **Voxel Size** | 4362 | **P-value** | 2.98E-07 |  |
| **Region** | **Side** | **X** | **Y** | **Z** | **Z-Score** |
| Anterior cingulate cortex | R | 6 | 16 | 32 | 6.55 |
| Anterior cingulate cortex | L | -2 | 12 | 30 | 5.53 |
| Supplemental motor area | R | 4 | 2 | 48 | 6.44 |
| Supplemental motor area | L | -2 | 6 | 48 | 5.21 |
| Midcingulate cortex | R | 2 | 16 | 24 | 5.77 |
| Midcingulate cortex | L | -2 | -14 | 30 | 4.97 |
| Superior frontal gyrus | R | 20 | 8 | 62 | 4.41 |
| Medial superior frontal gyrus | R | 2 | 42 | 48 | 3.43 |
| **Cluster 3** | **Voxel Size** | 1286 | **P-value** | 0.00207 |  |
| **Region** | **Side** | **X** | **Y** | **Z** | **Z-Score** |
| Cerebellum | L | -24 | -56 | -30 | 5.7 |
| **Deactivation** |  |  |  |  |  |
| **Cluster 1** | **Voxel Size** | 19085 | **P-value** | 7.33E-19 |  |
| **Region** | **Side** | **X** | **Y** | **Z** | **Z-Score** |
| Lateral occipital cortex | L | -36 | -82 | 32 | 6.38 |
| Lateral occipital cortex | R | 30 | -78 | 42 | 5.71 |
| Fusiform gyrus | R | 28 | -38 | -22 | 6.36 |
| Fusiform gyrus | L | -30 | -42 | -12 | 6.22 |
| Cerebellum | R | 30 | -46 | -24 | 5.89 |
| Precuneus | L | -8 | -54 | 18 | 5.86 |
| Precuneus | R | 8 | -58 | 20 | 5.71 |
| Hippocampus | L | -10 | -42 | 4 | 5.39 |
| Hippocampus | R | 30 | -28 | -10 | 3.73 |
| Lingual gyrus | R | 8 | -56 | -2 | 5.21 |
| Occipital Pole | L | -24 | -90 | 28 | 4.98 |
| Occipital Pole | R | 20 | -90 | 38 | 4.81 |
| Posterior cingulate cortex | L | -10 | -46 | 34 | 4.83 |
| Posterior cingulate cortex | R | 4 | -40 | 36 | 4.5 |
| Angular gyrus | L | -40 | -60 | 26 | 4.83 |
| Thalamus | L | -18 | -34 | 2 | 4.51 |
| Paracentral Lobule (M1/S1) | R | 4 | -36 | 60 | 4.3 |
| Paracentral Lobule (M1/S1) | L | -4 | -40 | 58 | 4.2 |
| Cuneus gyrus | R | 16 | -70 | 22 | 4.19 |
| Precentral gyrus | L | -26 | -20 | 58 | 3.89 |
| **Cluster 2** | **Voxel Size** | 680 | **P-value** | 0.0236 |  |
| **Region** | **Side** | **X** | **Y** | **Z** | **Z-Score** |
| Middle frontal gyrus | L | -26 | 24 | 34 | 5.05 |
| Superior frontal gyrus | L | -22 | 40 | 44 | 4.28 |

**Table S5. Comparison of brain responses to 2.5kg/cm^2^ between the adolescent group and the adult group (Z>3.1, p<0.05, cluster-corrected).**

| **Adolescents>Adults** |  |  |  |  |  |
| --- | --- | --- | --- | --- | --- |
| **Cluster 1** | **Voxel Size** | 625 | **P-value** | 9.14E-05 |  |
| **Region** | **Side** | **X** | **Y** | **Z** | **Z-Score** |
| Superior frontal gyrus | R | 20 | 60 | 34 | 4.29 |
| Medial superior frontal gyrus | R | 8 | 56 | 4 | 4.26 |
| Middle frontal gyrus | R | 44 | 48 | 26 | 4.02 |
| **Cluster 2** | **Voxel Size** | 230 | **P-value** | 0.0261 |  |
| **Region** | **Side** | **X** | **Y** | **Z** | **Z-Score** |
| Supramarginal gyrus | R | 68 | -34 | 38 | 5.23 |
| Angular gyrus | R | 66 | -48 | 22 | 3.81 |
| **Adolescents<Adults** |  |  |  |  |  |
| **Cluster 1** | **Voxel Size** | 430 | **P-value** | 0.00119 |  |
| **Region** | **Side** | **X** | **Y** | **Z** | **Z-Score** |
| Rectus gyrus | R | 10 | 32 | -18 | 4.49 |
| Rectus gyrus | L | -2 | 34 | -24 | 3.67 |
| Medial orbital gyrus | R | 16 | 42 | -20 | 4.23 |

**Table S6. Comparison of brain responses to 4kg/cm^2^ between the adolescent group and the adult group (Z>3.1, p<0.05, cluster-corrected).**

| **Adolescents>Adults** |  |  |  |  |  |
| --- | --- | --- | --- | --- | --- |
| **Cluster 1** | **Voxel Size** | 1152 | **P-value** | 4.83E-06 |  |
| **Region** | **Side** | **X** | **Y** | **Z** | **Z-Score** |
| Paracingulate | R | 4 | 48 | 26 | 4.96 |
| Medial superior frontal gyrus | R | 2 | 62 | 16 | 4.77 |
| Anterior cingulate cortex | R | 4 | 28 | 16 | 4.37 |
| Anterior cingulate cortex | L | -10 | 38 | 16 | 4.12 |
| Anterior cingulate cortex (rostral) | R | 4 | 38 | 10 | 3.97 |
| **Adolescents<Adults** |  |  |  |  |  |
| **Cluster 1** | **Voxel Size** | 250 | **P-value** | 0.0433 |  |
| **Region** | **Side** | **X** | **Y** | **Z** | **Z-Score** |
| Fusiform gyrus | L | -30 | -58 | -16 | 4.55 |
| Cerebellum | L | -22 | -50 | -22 | 4 |

**Table S7. Pain-evoked brain responses within brain networks (results of one-sample t-tests)**

| Stimulus Intensity: |  | 2.5kg/cm2 | | |  |  | 4kg/cm2 | |  |
| --- | --- | --- | --- | --- | --- | --- | --- | --- | --- |
| Group | | Adolescent | | Adult |  | Adolescent | | Adult |  |
| Brain Network | | t value | p value | t value | p value | t value | p value | t value | p value |
| Somatomotor | | 3.14 | 0.0038 | 6.17 | <0.0001 | 2.75 | 0.01 | 6.73 | <0.0001 |
| Dorsal Attentional | | -3.73 | 0.0008 | -2.2 | 0.0362 | -4.19 | 0.0002 | -2.04 | 0.05 |
| Ventral Attentional | | 13.07 | <0.0001 | 13.03 | <0.0001 | 16.4 | <0.0001 | 15.05 | <0.0001 |
| Frontoparietal | | 5.27 | <0.0001 | 4.64 | <0.0001 | 3.98 | 0.0004 | 2.5 | 0.0183 |
| Default Mode | | 1.55 | 0.1326 | -1.51 | 0.143 | 1.14 | 0.2627 | -3.32 | 0.0024 |
| Limbic | | -1.16 | 0.2548 | 1.32 | 0.1969 | 0.78 | 0.4427 | 0.83 | 0.4057 |
| Visual | | -5.13 | <0.0001 | -8.26 | <0.0001 | -7.24 | <0.0001 | 5.82 | <0.0001 |

**Table S8. Brain activity predictive of higher pain intensity ratings controlling for stimulus intensity in the adolescent group (Path b effect, q<0.05, FDR corrected, corresponding to p<0.000396)**

| **Region** | **Side** | **X** | **Y** | **Z** | **Z-Score** |
| --- | --- | --- | --- | --- | --- |
| Supplemental motor area | R | 12 | -10 | 72 | 5.88 |
| Fusiform gyrus | L | -26 | -10 | -40 | 5.13 |
| Superior temporal gyrus | L | -54 | -32 | 15 | 5 |
| Superior temporal gyrus | R | 60 | -32 | 15 | 4.74 |
| Middle cingulate cortex | R | 14 | -36 | 44 | 4.79 |
| Middle cingulate cortex | L | -10 | -36 | 48 | 4.44 |
| Cerebellum | L | -38 | -48 | -26 | 4.64 |
| Parahippocampal gyrus/ Hippocampus | L | -28 | -6 | -26 | 4.62 |
| Parahippocampal gyrus/ Hippocampus | R | 22 | -2 | -22 | 4.35 |
| Amygdala/Hippocampus | R | 24 | -6 | -18 | 4.14 |
| Superior parietal lobule | L | -20 | -50 | 68 | 4.47 |
| Precentral gyrus (M1) | R | 46 | -4 | 60 | 4.44 |
| Precentral gyrus (M1) | L | -32 | -20 | 51 | 3.64 |
| Middle temporal gyrus | L | -48 | -68 | 8 | 4.41 |
| Middle temporal gyrus | R | 50 | -66 | 6 | 3.66 |
| Parietal Operculum (S2) | R | 50 | -22 | 15 | 4.24 |
| Posterior insular cortex | R | 36 | -18 | 14 | 4.2 |
| Postcentral gyrus (S1) | L | -22 | -36 | 67 | 4.11 |
| Inferior temporal gyrus | L | -56 | -50 | -20 | 4.05 |
| Inferior temporal gyrus | R | 53 | -32 | -25 | 3.76 |
| Superior frontal gyrus (DLPFC) | L | -23 | -6 | 62 | 3.94 |
| Middle frontal gyrus (DLPFC) | L | -40 | 43 | 32 | 3.85 |
| Paracentral lobule | R | 2 | -26 | 68 | 3.68 |
| Putamen | R | 32 | -12 | 2 | 3.52 |

**Table S9. Brain activity predictive of higher pain intensity ratings controlling for stimulus intensity in the adult group (Path b effect, q<0.05, FDR corrected, corresponding to p<0.0017)**

| **Region** | **Side** | **X** | **Y** | **Z** | **Z-Score** |
| --- | --- | --- | --- | --- | --- |
| Fusiform gyrus | R | 26 | -8 | -46 | 6.28 |
| Midcingulate cortex | R | 2 | -10 | 42 | 4.66 |
| Posterior insula/ central operculum | R | 46 | -12 | 14 | 4.53 |
| Middle frontal gyrus | L | -36 | 60 | 4 | 4.41 |
| Middle temporal gyrus | L | -65 | -28 | -14 | 4.33 |
| Middle temporal gyrus | R | 44 | -58 | 8 | 3.32 |
| Cerebellum | R | 24 | -52 | -60 | 4.28 |
| Cerebellum | L | -36 | -66 | -58 | 3.73 |
| Supplemental motor area | R | 8 | -12 | 76 | 4.22 |
| Paracentral lobule | R | 6 | -40 | 78 | 4.08 |
| Thalamus | L | -16 | -20 | -2 | 4.02 |
| Lingual gyrus | L | -12 | -88 | -14 | 3.98 |
| Lingual gyrus | R | 11 | -52 | -4 | 3.93 |
| Cuneus | L | -8 | -88 | 42 | 3.93 |
| Superior temporal gyrus | R | 60 | -8 | 0 | 3.93 |
| Superior temporal gyrus | L | -50 | -12 | 0 | 3.65 |
| Globus pallidum | R | 16 | 0 | 6 | 3.87 |
| Globus pallidum | L | -20 | 0 | -4 | 3.4 |
| Superior medial frontal gyrus (mPFC) | R | 4 | 69 | 8 | 3.78 |
| Inferior temporal gyrus | R | 54 | -19 | -30 | 3.44 |
| Parietal Operculum (S2) | R | 52 | -20 | 16 | 3.44 |
| Superior parietal lobule | L | -36 | -55 | 60 | 3.43 |
| Postcentral gyrus (S1) | R | 40 | -32 | 48 | 3.28 |

**Table S10. Brain activity predictive of higher pain intensity ratings controlling for stimulus intensity greater in adolescents than in adults (Moderated Path b effect, uncorrected p<0.001)**

| **Region** | **Side** | **X** | **Y** | **Z** | **Z-Score** |
| --- | --- | --- | --- | --- | --- |
| Middle temporal gyrus | R | 64 | -46 | 10 | 3.71 |
| Middle temporal gyrus | L | -46 | -68 | 10 | 3.64 |
| Superior frontal gyrus (Premotor cortex) | L | -22 | -7 | 60 | 3.63 |
| Superior frontal gyrus | R | 27 | 10 | 61 | 3.33 |
| Hippocampus/Amygdala | L | -37 | -4 | -26 | 3.61 |
| Hippocampus/Amygdala | R | 32 | -8 | -15 | 3.45 |
| Parahippocampal gyrus | R | 30 | 2 | -30 | 3.58 |
| Parahippocampal gyrus | L | -20 | 2 | -32 | 3.48 |
| Superior orbital frontal gyrus | R | 11 | 18 | -19 | 3.58 |
| Middle cingulate cortex | R | 12 | -38 | 48 | 3.48 |
| Paracentral lobule (S1) | R | 4 | -34 | 58 | 3.40 |
| Postcentral gyrus (S1) | L | -19 | -33 | 70 | 3.37 |
| Precentral gyrus (M1) | R | 22 | -18 | 61 | 3.24 |

**Table S11. Brain activity mediating the relationship between stimulus intensity and pain intensity ratings in the adolescent group (Path ab effect, q<0.05, FDR corrected, corresponding to p<0.000396)**

| **Region** | **Side** | **X** | **Y** | **Z** | **Z-Score** |
| --- | --- | --- | --- | --- | --- |
| Amygdala | L | -24 | -2 | 25 | 5.6 |
| Amygdala | R | 33 | 0 | -22 | 3.78 |
| Parahippocampus | R | 24 | -26 | -20 | 4.76 |
| Parahippocampus | L | -24 | 2 | -33 | 3.44 |
| Middle frontal gyrus (DLPFC) | R | 36 | 30 | 32 | 4.75 |
| Inferior frontal gyrus (DLPFC) | L | -48 | 21 | 24 | 4.43 |
| Cerebellum | R | 11 | -54 | -8 | 4.47 |
| Cerebellum | L | -29 | -58 | -24 | 3.6 |
| Middle temporal gyrus | R | 52 | -30 | -14 | 4.38 |
| Supramarginal gyrus | R | 48 | -38 | 38 | 4.29 |
| Putamen/Globus Pallidum | L | -22 | 0 | -6 | 4.12 |
| Putamen | R | 19 | 2 | 0 | 3.59 |
| Middle cingulate cortex | R | 6 | 30 | 36 | 3.88 |
| Middle cingulate cortex | L | -8 | 6 | 32 | 3.78 |
| Anterior cingulate cortex | R | 10 | 20 | 28 | 3.84 |
| Superior medial frontal gyrus (mPFC) | R | 4 | 63 | 8 | 3.7 |
| Inferior temporal gyrus | L | -46 | 8 | -41 | 3.64 |
| Inferior temporal gyrus | R | 44 | -30 | -16 | 3.48 |
| Middle frontal gyrus (VLPFC) | L | 41 | 48 | 16 | 3.61 |

**Table S12. Brain activity mediating the relationship between stimulus intensity and pain intensity ratings greater in the adult group (Path ab effect, q<0.05, FDR corrected, corresponding to p<0.0017)**

| **Region** | **Side** | **X** | **Y** | **Z** | **Z-Score** |
| --- | --- | --- | --- | --- | --- |
| Inferior temporal gyrus | L | -42 | -2 | -32 | 5.93 |
| Cerebellum | R | 38 | -70 | -34 | 5.66 |
| Cerebellum | L | -26 | -56 | -56 | 4.16 |
| Superior temporal gyrus | R | 48 | -18 | 10 | 5.5 |
| Superior temporal gyrus | L | -56 | 0 | -14 | 4.89 |
| Postcentral gyrus | L | -64 | -12 | 30 | 5.35 |
| Precentral gyrus | L | -26 | 58 | 49 | 5 |
| Middle frontal gyrus (DLPFC) | R | 27 | 26 | 36 | 4.84 |
| Middle frontal gyrus (DLPFC) | L | -28 | 24 | 50 | 4.83 |
| Supplemental motor area | R | 2 | 2 | 72 | 4.82 |
| Supplemental motor area | L | -2 | -4 | 64 | 4.5 |
| Paracentral lobule | L | -4 | -34 | 68 | 4.77 |
| Supramarginal gyrus | R | 52 | -14 | 26 | 4.63 |
| Globus pallidum | R | 14 | 2 | -4 | 4.36 |
| Anterior cingulate cortex | L | -10 | 34 | 30 | 4.35 |
| Anterior cingulate cortex | R | 2 | 22 | 18 | 4.18 |
| Amygdala | L | -26 | -4 | -11 | 4.29 |
| Middle cingulate cortex | L | -16 | -27 | 56 | 4.2 |
| Middle cingulate cortex | R | 2 | 2 | 40 | 3.74 |
| Middle frontal gyrus (VLPFC) | R | 42 | 29 | 12 | 4.04 |
| Middle frontal gyrus (VLPFC) | R | 42 | 55 | 4 | 3.91 |

**Table S13. Brain activity mediating the relationship between stimulus intensity and pain intensity ratings greater in adolescents than in adults (Moderated Path ab effect, uncorrected p<0.001)**

| **Region** | **Side** | **X** | **Y** | **Z** | **Z-Score** |
| --- | --- | --- | --- | --- | --- |
| Inferior temporal gyrus | R | 55 | -32 | -14 | 6 |
| Inferior temporal gyrus | L | -41 | 9 | -41 | 4.15 |
| Middle frontal gyrus (VLPFC) | R | 40 | 44 | 8 | 5.31 |
| Superior temporal gyrus | R | 46 | -20 | -3 | 5.8 |
| Superior medial frontal gyrus (mPFC) | R | 5 | 55 | 20 | 5.22 |
| Superior medial frontal gyrus (mPFC) | R | 4 | 43 | 43 | 3.57 |
| Anterior cingulate cortex | R | 9 | 22 | 27 | 3.89 |
| Superior frontal gyrus (DLPFC) | R | 20 | 29 | 49 | 3.74 |
| Middle cingulate cortex | R | 5 | 29 | 34 | 3.59 |
| Middle insula cortex | L | -40 | 9 | -8 | 3.57 |
| Inferior frontal gyrus (DLPFC) | L | -46 | 14 | 29 | 3.74 |
| Inferior frontal gyrus (DLPFC) | R | 43 | 13 | 28 | 3.18 |
| Pararhippocampus gyrus | R | 26 | -12 | -25 | 3.31 |
